# Loss of the tumor suppressor NUMB drives aggressive bladder cancer through hyperactivation of a RhoA/ROCK/YAP signaling circuitry

**DOI:** 10.1101/2024.08.27.609735

**Authors:** F. A. Tucci, R. Pennisi, D. C. Rigiracciolo, M. G. Filippone, R. Bonfanti, F. Romeo, S. Freddi, C. Soriani, S. Rodighiero, G. Jodice, F. Sanguedolce, G. Renne, N. Fusco, P.P. Di Fiore, G. Pruneri, G. Musi, G. Vago, D. Tosoni, S. Pece

## Abstract

Bladder cancer (BCa) is one of the most challenging and costly cancers to treat, yet little progress has been made on the development of predictive biomarkers and targeted therapies. Here, we uncover a critical function of Numb as a tumor suppressor in the bladder, identifying loss of Numb expression as a causal alteration in BCa that underlies biological aggressiveness and disease progression. Through retrospective cohort studies, we established that a Numb-deficient tumor status correlates with worse overall survival in post-cystectomy muscle-invasive bladder cancer (MIBC) patients and increased risk of MIBC progression in non-muscle-invasive bladder cancer (NMIBC) patients. The prognostic value of Numb loss can be attributed to its crucial role as a determinant of aggressive bladder tumorigenesis, as demonstrated in mouse and human models. Targeted *Numb* ablation in the basal layer of the urothelium was alone sufficient to trigger spontaneous bladder tumorigenesis and drive progression from preneoplastic to preinvasive and, ultimately, overtly invasive tumors. Additionally, *Numb* ablation sensitized the urothelium to other oncogenic insults, accelerating tumor onset and progression. Using 3D-Matrigel organoid cultures to recapitulate bladder tumorigenesis *in vitro*, we found that Numb loss heightens the proliferative and invasive potential of both mouse and human BCa cells. Integrative transcriptomic and functional analyses revealed that downregulation of the canonical Hippo pathway, resulting in enhanced YAP transcriptional activity, underlies the biological aggressiveness of Numb-deficient BCa. These molecular events are dependent on the activation of RhoA/ROCK signaling subsequent to Numb loss. Thus, a dysfunctional Numb–RhoA/ROCK–Hippo/YAP regulatory network is at play in aggressive Numb-deficient BCa and represents a therapeutic vulnerability. A 27-gene prognostic signature capable of identifying high-risk Numb-deficient patients could provide the basis of a clinical tool to stratify patients for innovative RhoA/ROCK/YAP targeted therapies.

**One Sentence Summary:** Numb loss-directed hyperactivation of RhoA/ROCK/YAP underlies aggressive bladder cancer biology.

## INTRODUCTION

Bladder cancer (BCa) ranks among the most common neoplasms in industrialized countries^1^. Most BCa manifest as urothelial carcinomas, originating from the transitional epithelium lining the inner surface of the bladder^2–4^. Clinical staging of these tumors depends on the extent of invasion into the bladder wall and their grade of cellular differentiation^2,4^. Tumors that have penetrated the superficial layers, infiltrating the detrusor muscle and beyond, are classified as muscle-invasive BCa (MIBC), and constitute ∼25% of all BCa. Despite aggressive treatments, such as radical cystectomy and adjuvant platinum-based chemotherapy, MIBC has an overall poor prognosis, characterized by a high metastatic potential and a 5-year overall survival of ∼50%. The aggressive treatments are also associated with high morbidity, severe side effects and a diminished quality of life^4–6^.

Conversely, tumors confined to the inner lining of the bladder are classified as non-muscle invasive BCa (NMIBC), representing ∼75% of newly diagnosed BCa cases^2,7^. NMIBCs are highly heterogeneous, comprising tumors of different stage and grade: i) Ta papillary tumors, confined to the mucosa and typically low-grade/well-differentiated; ii) T1 tumors with submucosal invasion, representing a heterogeneous group that can present as either low-or high-grade/poorly differentiated lesions; and iii) carcinoma *in situ* (CIS; Tis in the TNM classification), typically presenting as flat and high-grade dysplasia lesions^4^. Although 5-year survival rates for NMIBC are favorable (>90%), a significant proportion of patients experience disease recurrence (∼50-70%) and progression to MIBC disease (∼20-30%)^8^. Notably, NMIBC patients who progress to MIBC often face a worse prognosis compared to patients with primary MIBC^9^. This, coupled with the lack of improvement in MIBC mortality rates, makes the transition from NMIBC to MIBC a life-threatening event. Therefore, high-risk NMIBC patients must undergo lifelong cystoscopic surveillance with bladder-sparing transurethral resection (TUR), often followed by adjuvant intravesical instillations of Bacillus Calmette-Guérin (BCG) or bladder perfusion chemotherapy to eradicate residual disease and reduce the frequency of recurrence and progression^4,6,10^. This treatment regimen makes BCa the most expensive cancer to treat^11^.

The considerable heterogeneity of NMIBC presents challenges in its clinical management, exacerbated by the inadequacy of current staging criteria which heavily rely on clinicopathological characteristics. Incorrect staging can lead to understaging and subsequent undertreatment, negatively affecting survival, or overtreatment with early cystectomy, associated with significant morbidity^8^. Moreover, despite the numerous molecular alterations described for this disease, targeted therapies to prevent disease progression are currently lacking^2,12–15^. Consequently, there is an urgent need for biomarkers of disease progression to guide clinical decision-making, particularly regarding the choice between conservative (surveillance and TUR) *vs.* aggressive treatments (early cystectomy plus adjuvant therapies). A deeper understanding of the biology underlying the NMIBC to MIBC transition is therefore essential for an improved personalized management of NMIBC patients^3,8,10,13^.

In this study, we have elucidated a critical role of the loss of the tumor suppressor Numb in bladder tumorigenesis and disease progression. Through retrospective cohort studies, we demonstrate that a Numb-deficient status is a prognostic biomarker for risk of MIBC progression in NMIBC patients, and for poor prognosis in MIBC patients. Furthermore, using a Numb-knockout (KO) mouse model, we establish that *Numb* ablation is alone sufficient to drive the stepwise malignant transformation of the urothelium, causing the appearance of preneoplastic lesions, preinvasive tumors, and muscle-invasive tumors. Integrating functional and molecular studies in mouse and human BCa models, we show that Numb loss leads to the upregulation of RhoA/ROCK signaling to the actin cytoskeleton. This, in turn, results in the downregulation of the Yes-associated protein-1 (YAP1) inhibitory Hippo pathway, leading to hyperactivation of YAP transcriptional activity. Through genetic and pharmacological inhibition studies in mouse and human BCa models, we establish that hyperactivation of RhoA/ROCK/YAP signaling is responsible for the aggressive migratory/invasive phenotype of Numb-deficient BCa cells. These findings highlight the RhoA/ROCK/YAP circuitry as a therapeutic vulnerability in Numb-deficient NMIBC patients that can be targeted pharmacologically to prevent progression to MIBC.

## RESULTS

### Numb loss is a hallmark of aggressive disease in human BCa patients

A preliminary immunohistochemistry (IHC) analysis of formalin-fixed paraffin-embedded (FFPE) BCa samples obtained from patients who underwent radical cystectomy, revealed that Numb expression is frequently downregulated in the tumor lesion compared to the adjacent normal urothelium (**Fig. 1a**). This observation prompted the investigation of the relevance of Numb loss to the natural history of the human BCa. We first surveyed a retrospective consecutive cohort of 356 MIBC patients subjected to radical cystectomy, assessing Numb status by IHC on whole FFPE tumor sections. Complete clinicopathological follow-up (median, 5.73 years) was available for 258 patients. Approximately 80% of patients were categorized as Numb^High^ and ∼20% as Numb^Low^. Notably, a Numb^Low^ status significantly correlated with clinical parameters of aggressive disease (e.g., tumor extension and vascular invasion) and positive lymph node status, and also predicted a higher rate of mortality, independently of standard clinicopathological risk factors (Numb^Low^ *vs.* Numb^High^, HR=1.64, CI=1.1-2.4, p=0.0098) (**Fig. 1b** and **Extended Data Fig. 1a,b**).

**Figure 1.**
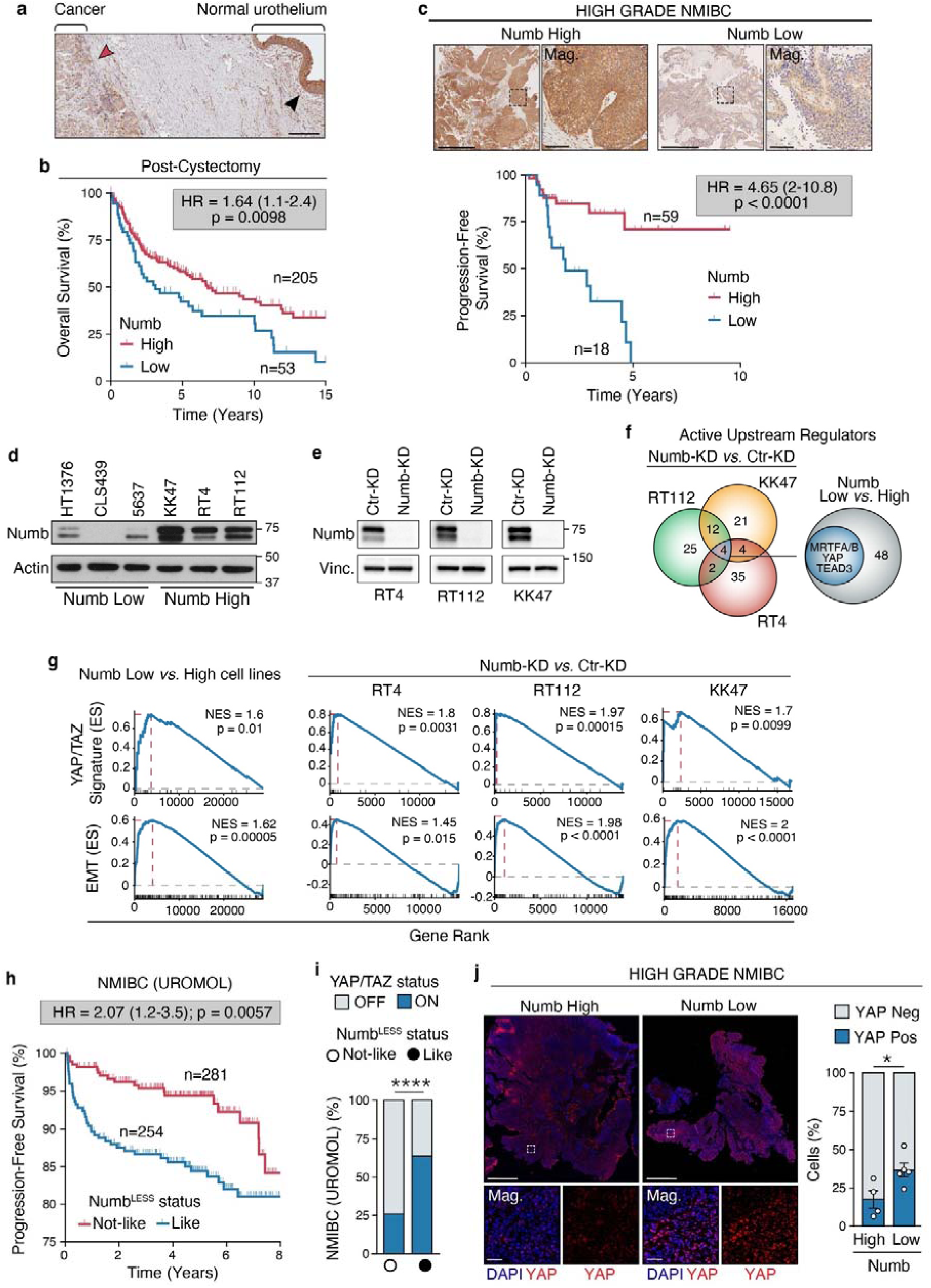
Numb loss is prognostic in human BCs and correlates with NMIBC progression and YAP activation. **a.** Representative IHC image of an area of Numb-deficient MIBC (red arrowhead) bordering the normal urothelium expressing Numb (black arrowhead). Bar, 400 μm. **b**. Overall survival of post-cystectomy MIBC patients (n=258) stratified by Numb expression (Numb^High^ *vs.* Numb^Low^) determined by IHC (see also Extended Data Fig. 1a and 1b for cohort description and multivariable analysis). HR, hazard ratio (95% confidence interval); p, Log Rank test p-value; n, patient number; in this and all other relevant panels. **c.** Top, Representative IHC images of Numb expression in high-grade Numb^High^ and Numb^Low^ NMIBC. Boxed areas are magnified (Mag.) on the right. Bars, 1 mm; Mag. 100 µm. Bottom, progression-free survival of NMIBC patients (n=77) stratified by Numb expression (see also Extended Data Fig. 1c and 1d for cohort description and multivariable analysis). **d**. Immunoblot of Numb expression in MIBC Numb^Low^ cell lines (HT1376, CLS439, 5637) and NMIBC Numb^High^ cell lines (KK47, RT4, RT112). Actin, loading control. **e.** Immunoblot of Numb expression in RT4, RT112 and KK47 NIMBC cells knocked down for Numb by siRNA (Numb-KD) or mock siRNA in controls (Ctr-KD). Vinculin (Vinc.), loading control. **f.** The Venn diagram shows the number of activated transcriptional regulators and their intersection identified by the independent comparison of the RT4, RT112 and KK47 NIMBC cells, Numb-KD *vs*. Ctr-KD. The 4 common YAP/TAZ transcriptional regulators (MRTFA/B, YAP and TEAD3) appearing in the Venn diagram intersection on the left were among the 52 activated transcriptional regulators identified in the integrative transcriptomic analysis of the Numb^Low^ *vs*. Numb^High^ cell lines, as indicated in the blue circle (see Extended Data Fig. 2a for the complete list of regulators). **g.** GSEA showing enrichment of an active YAP/TAZ (top panels) and EMT (bottom panels) gene signature in the set of common differentially expressed genes across the 3 NUMB^Low^ *vs.* 3 Numb^High^ cell lines (left) and between the Numb-KD *vs*. CTR-KD condition in each of the indicated Numb^High^ cell lines (right). n[=[2 for each condition of each cell line; ES, Enrichment Score; NES, Normalized Enrichment Score; p, permutation test p-value. **h.** Progression-free survival of NMIBC patients of the UROMOL cohort (n=535) stratified with the Numb^LESS^ signature into Numb^LESS^-Like (n=254) or NUMB^LESS^-Not-Like (n=281) groups, corresponding to a Numb-deficient or Numb-proficient status, respectively (see also Extended Data Fig. 2d for multivariable analysis). **i.** NMIBC patients of the UROMOL cohort stratified with the Numb^LESS^ signature into Numb^LESS^-Like or NUMB^LESS^-Not-Like groups (as shown in h) were further classified as YAP/TAZ active (ON) or inactive (OFF) based on the expression of the 22-gene YAP/TAZ signature (as in g) (see also Extended Data Fig. 2e and Methods). ****, p<0.0001, by Fisher’s exact test. **j.** Left, Immunofluorescence analysis of YAP expression (red) and DAPI nuclear stain (blue) in high-grade Numb^High^ *vs.* Numb^Low^ NMIBC tumors. Boxed areas in the upper panels are magnified (Mag.) in the lower panels. Bars, 1 mm in upper panels; Mag., 50 μm. Right, quantification of the % of YAP-positive cells expressed as mean ± SEM of 4 Numb^High^ and 5 Numb^Low^ NIMBC. *, p=0.038, by Welch’s *t*-test.

To investigate whether Numb expression also has clinical value in NMIBC, we analyzed a cohort of 77 high-grade NMIBC patients who progressed or not to MIBC in a 4-month minimum follow-up period, as assessed by TUR (median follow-up 1.94 years; see **Extended Data Fig. 1c** for clinicopathological characteristics). Strikingly, in this cohort, a Numb^Low^ status predicted a worse progression-free survival with a higher rate of MIBC progression compared to Numb^High^ patients (HR=4.65, CI=2 – 10.8, p<0.0001) (**Fig. 1c**). These retrospective clinical studies clearly indicate that a Numb-deficient status is a hallmark of aggressive disease and poor prognosis in BCa. In NMIBC patients, Numb status behaves as a predictive biomarker of risk of muscle-invasion progression independently of other clinical parameters (**Extended Data Fig. 1d**). Thus, Numb status could guide clinical decision-making between conservative *vs.* more aggressive therapies.

### Numb loss results in pathological regulation of YAP transcriptional activity in BCa

To investigate transcriptional programs associated with Numb loss in human BCa, we conducted a comparative analysis of the transcriptomes (obtained by bulk RNA-seq) of paired Numb-deficient and Numb-proficient BCa model systems. Specifically, we compared three MIBC Numb^Low^ cell lines (CLS439, 5637 and HT1376) with three NMIBC Numb^High^ cell lines (RT4, RT112 and KK47) (**Fig. 1d**). Additionally, for an isogeneic comparison, we examined the three NMIBC Numb^High^ cell lines silenced or not for Numb (Numb-KD *vs.* Ctr-KD) (**Fig. 1e**). Integrative analysis of transcriptomic data using the IPA ‘Upstream Regulator Analysis’, led to the identification of common transcriptional regulators that are upregulated in the Numb-deficient condition relative to the Numb-proficient condition (i.e., in NUMB^Low^/Numb-KD *vs.* NUMB^High^/Ctr-KD, respectively). The most distinctive molecular traits associated with the Numb-deficient condition were activation of upstream regulators of the YAP/TAZ pathway (e.g., MRTFA/B, TEAD3 and YAP) and the EMT/phenotypic plasticity pathways (e.g., CTNNB1, KLF4, TWIST1, SOX4, SNAI, ZEB1) (**Fig. 1f** and **Extended Data Fig. 2a**). In line with these results, GSEA analysis of the set of differentially expressed genes between the paired Numb-deficient *vs*. Numb-proficient cell model systems revealed the enrichment of a YAP/TAZ signature, composed of 22 transcriptional targets of the pathway^16^, and an EMT signature (“Cancer Hallmarks”, see Materials and Methods), in the Numb-deficient condition (**Fig. 1g**).

### Derivation of a clinically relevant gene signature characteristic of the Numb-deficient state

Previous studies in breast cancer have shown that, beyond the overall decrease in total Numb protein levels^17^, other molecular events, such as Numb isoform-specific transcriptional regulation and post-translational modifications, may underlie a deficient Numb status and correlate with clinical aggressiveness^18,19^. Considering the possibility of a similar scenario in BCa, we reasoned that a gene signature representative of the pattern of molecular alterations associated with a Numb-deficient state could be a more precise tool, compared to the direct assessment of total Numb IHC levels, to identify patients that will progress from NMIBC to MIBC. With this idea in mind, we performed an integrative comparative analysis of the transcriptomes of Numb-KD *vs.* Ctr-KD RT4, RT112 and KK47 NMIBC cell lines to identify the most significantly deregulated genes (absolute Log_2_ fold-change>1, FDR adjusted p<0.01) displaying a consistent direction of regulation (up or down) relative to Numb status across the three paired analyses. This analysis led to the identification of a minimal signature composed of 27 genes up-or down-regulated consequent to Numb-KD in the three NMIBC cell lines, characteristic of the Numb defective condition (referred to hereafter as the “Numb^LESS^ signature”). In an unsupervised clustering analysis, this signature correctly stratified the Numb^High^ (RT4, RT112 and KK47) and Numb^Low^ (CLS439, 1537 and HT1376) BCa cell lines into two distinct groups **(Extended Data Fig. 2b).**

To test the clinical value of the Numb^LESS^ signature in NMIBC patients, we interrogated the transcriptomes of 535 NMIBC patients, with complete long-term clinicopathological follow-up (median follow-up, 5.12 months), who were included in the UROMOL study^20^. The unsupervised clustering analysis of this cohort using the Numb^LESS^ signature identified two distinct groups of NMIBC patients who were classified as Numb^LESS^*-*like (corresponding to a Numb-deficient status) or Numb^LESS^*-*not-like (corresponding to a Numb-proficient status) (**Extended Data Fig. 2c**). Numb^LESS^*-*like NMIBC patients displayed a significantly higher risk of MIBC progression and shorter progression-free survival compared to Numb^LESS^*-*not-like patients (HR=2.07, CI=1.2-3.5, p=0.0057) (**Fig. 1h**). Remarkably, the ability of the Numb^LESS^ signature to predict risk of MIBC progression was maintained in a multivariable analysis adjusted for other clinicopathological factors (**Extended Data Fig. 2d**). This finding, together with the results from the retrospective analysis of the NIMBC cohort by Numb IHC (see **Fig. 1c**), strongly argue for the clinical value of assessing Numb status in NMIBC patients to predict the risk of MIBC progression, beyond currently available staging parameters. It is also worthy of note that, in keeping with our initial hypothesis, the Numb^LESS^ 27-gene signature appears to have a superior stratification power, compared to IHC, for the identification of high-risk Numb-deficient NMIBC patients (∼50% Numb^LESS^*-*like *vs.* 20% IHC Numb^Low^), likely due to its ability to capture a dysfunctional Numb status associated with heterogeneous molecular events^18,19^, other than the Numb protein hyperdegradation initially described in breast cancer^17^.

### Relevance of YAP/TAZ pathway activation in BCa patients

Having established a link between Numb loss and YAP/TAZ pathway activation in mouse and human BCa cell models, we extended our investigation to examine the relevance of this signaling pathway to NMIBC patients. Notably, in the UROMOL cohort, we observed a significant enrichment of a 22-gene YAP/TAZ signature^16^ in the transcriptomic profiles of Numb^LESS^-like tumors, compared with Numb^LESS^*-*not-like tumors (**Fig. 1i and Extended Data Fig. 2e**). Consistently, by immunofluorescence (IF) analysis of high-grade NMIBC TUR samples, we observed a direct correlation between low expression of Numb protein and increased nuclear accumulation of YAP, indicative of activated YAP signaling^21^ (**Fig. 1j**). These results argue that the association between a Numb-deficient status and YAP/TAZ pathway activation observed in cell lines might be pathogenetically relevant to NMIBC patients.

### Numb deficiency drives malignant transformation of the mouse urothelium and accelerates carcinogen-induced bladder tumorigenesis

To investigate the impact of Numb dysfunction on homeostasis of the urothelium, we used a Numb knockout (Numb-KO) mouse model bearing targeted deletion of the *Numb* gene in the basal CK5+ layer (**Fig. 2a**). The Numb-KO mouse was generated by crossing the Cre-loxP conditional Numb*-* KO mouse (Numb^lox/stop/lox^) with the CK5-Cre mouse as previously described^22,23^. The rationale of this experimental strategy was based on evidence that, in both the mouse and human normal urothelium, Numb is more highly expressed in basal (CK5+/CK14+) cells compared to suprabasal intermediate (CK7+) cells and superficial umbrella (CK20+) cells (**Extended Data Fig. 3a,b**). We also reasoned that this model could have the potential to highlight key features of disease progression, considering previous evidence implicating basal, but not suprabasal, cells as the population-of-origin of aggressive BCa^24–26^.

**Figure 2.**
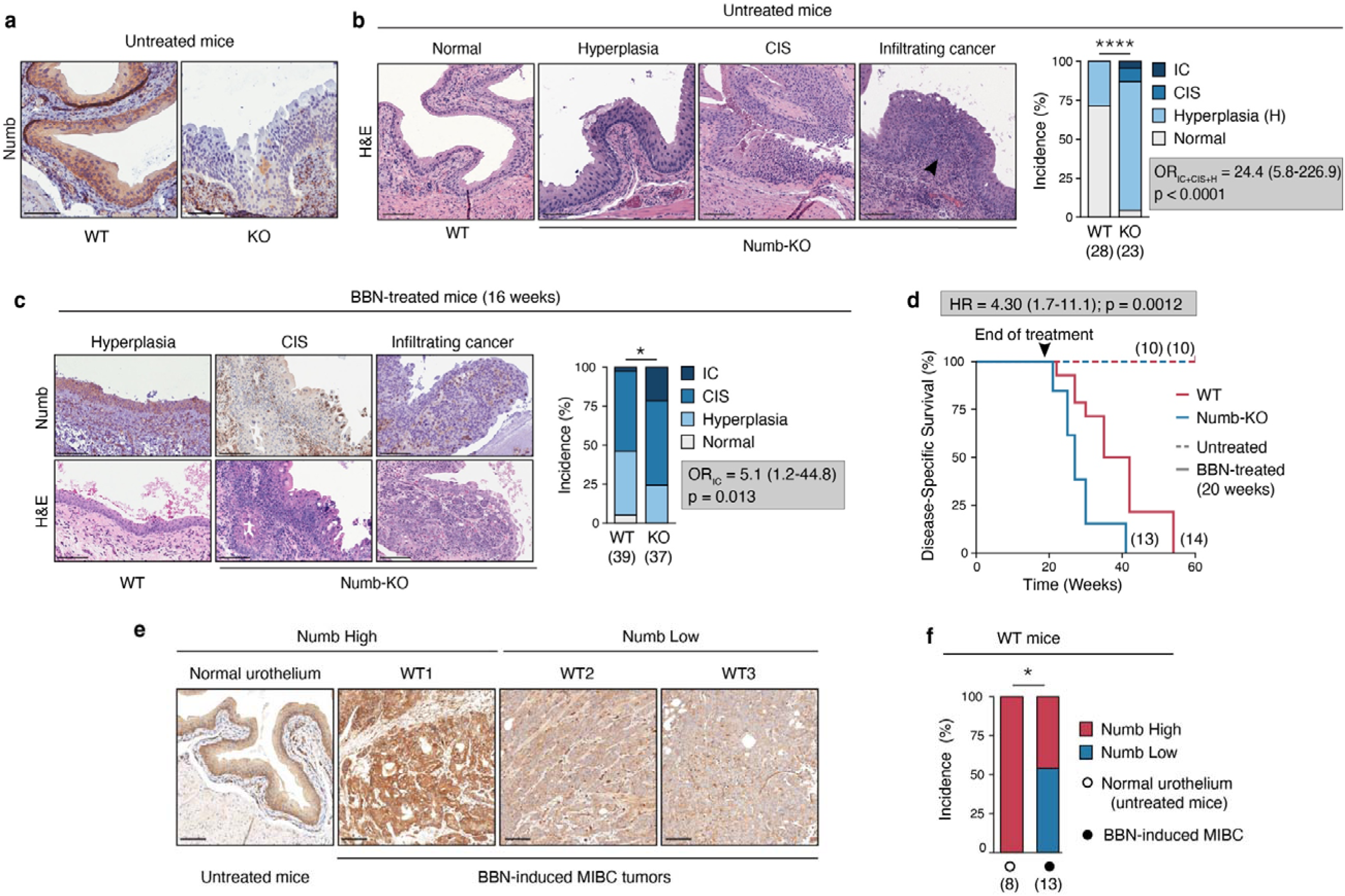
Numb loss drives spontaneous malignant transformation of the normal urothelium and accelerates carcinogen-induced bladder tumorigenesis. **a.** Representative IHC images showing endogenous Numb expression in the urothelium of untreated adult WT (WT) and Numb-KO (KO) mice. Bars, 100 μm. **b.** Left, Representative H&E images of the normal WT urothelium, and preneoplastic (Hyperplasia) and neoplastic (carcinoma *in situ*, CIS; invasive cancer, IC) urothelial lesions from Numb-KO mice. The black arrowhead indicates IC. Bars, 100 μm. Right, quantification of the incidence (%) of histological phenotypes in the urothelium of untreated, aged-matched 4-to 12-month old WT (n=28) and Numb-KO (n=23) mice. ****, p=0.00002 by Pearson’s Chi-squared test. Odds ratio (OR) with 95% CI and associated p-value by Fisher’s exact test is shown. **c.** Aged-matched 8-to 16-week-old WT (n=39) and Numb-KO (n=37) mice were exposed to 0.05% BBN in the drinking water for 16 weeks followed by a 2-week washout period (see Extended Data Fig. 3d). Bladder tissues were harvested and examined for histological changes (H&E) and Numb expression (IHC). Left, Representative images of a preneoplastic (Hyperplasia) lesion from BBN-WT mice and neoplastic (CIS and IC) lesions from BBN-Numb-KO mice. Right, quantification of the incidence (%) of histological phenotypes in WT *vs.* KO mice at the end of treatment. Bars, 100 μm. *, p=0.025 by Pearson’s Chi-squared test; OR (95% CI) with associated p-value by Fisher’s exact test is shown. **d.** Kaplan-Meier plot showing the disease-specific survival (%) of aged-matched 8-to 16-week-old WT (n=14) and Numb-KO (n=13) mice treated for 20 weeks with BBN before switching to regular drinking water. Mice were sacrificed according to endpoints (see Extended Data Fig. 3d and Methods). Dashed-line, WT (n=10) and Numb-KO (n=10) untreated mice. HR, (95% CI) with p-value by Log Rank test are shown. **e.** MIBC lesions excised from BBN-WT mice (n=13) were compared to the normal urothelium of untreated WT mice (n=8) for Numb expression by IHC. Shown are three MIBC lesions, one Numb^High^ (WT1) and two Numb^Low^ (WT2, WT3), and the normal Numb^High^ urothelium of untreated mice. Bars, 100 μm. **f.** Quantification of the experiment in ‘e’ showing the incidence (%) of Numb^Low^ *vs.* Numb^High^ MIBC tumors induced by BBN treatment in WT mice. *, p=0.018 by Fisher’s exact test.

Comparative histology of the Numb-KO *vs*. WT bladder mucosa revealed that the absence of Numb in the urothelium leads to its early hyperplastic thickening, characterized by expansion of the basal layer, and appearance of *in situ* (CIS) and overtly infiltrating neoplastic lesions (**Fig. 2b** and **Extended Data Fig. 3c**). These results suggest a direct involvement of Numb loss in initiating early morphological alterations in the urothelium, occurring prior to the emergence of preneoplastic lesions with a marked propensity to progress to advanced tumor stages.

To investigate the potential cooperative effect of Numb loss in bladder tumorigenesis, beyond its primary role, we examined the susceptibility of Numb-KO *vs*. WT mice to prolonged exposure to the chemical carcinogen, N-butyl-N-(4-hydroxybutyl) nitrosamine (BBN) (**Extended Data Fig. 3d**). Of note, the BBN-induced BCa mouse model faithfully reproduces the histological features and genetic alterations observed in human BCa, including the progression from NMIBC to MIBC^25,27,28^. Numb-KO mice displayed increased susceptibility to BBN, evidenced by a dramatically increased incidence and accelerated rate of formation of invasive carcinoma, compared to WT mice (OR=5.1, CI=1.2-44.8, p=0.013) (**Fig. 2c**), and a significant reduction in survival rate (HR=4.30, CI=1.7-11.1, p=0.0012) (**Fig. 2d**). Remarkably, we also noted that ∼50% of BBN-induced invasive tumors that developed in WT mice exhibited spontaneous loss of Numb expression (**Fig. 2e,f**).

Together, these results highlight the potent pro-tumorigenic effects of Numb loss in the urothelium: not only is Numb loss *per se* sufficient to induce bladder tumorigenesis, it also expedites invasive tumorigenesis driven by other oncogenic insults. Additionally, it is a frequently selected event during the neoplastic transformation of the urothelium, presumably as a means to alleviate its tumor suppressor function.

### Loss of Numb induces aggressive biological phenotypes in BCa cells through YAP transcriptional hyperactivation

Based on the above results, we reasoned that the paired WT *vs.* Numb-KO and BBN-WT *vs.* BBN-Numb-KO mouse models were the ideal preclinical setting to elucidate the molecular mechanisms underlying the aggressive biological behavior of Numb-deficient BCa. We focused our attention on the link between deficient Numb expression and YAP/TAZ pathway hyperactivation, given evidence of this interaction in human BCa cell lines and the enrichment in YAP/TAZ transcriptional targets observed in the prognostic Numb^LESS^-signature.

IHC analysis of endogenous YAP expression in the normal human and murine urothelium showed higher levels of YAP in the basal layer, with an evident nuclear staining (**Extended Data Fig. 4a**). Further analysis comparing the urothelium of WT *vs.* Numb-KO mice by *in situ* IF with an antibody specific to active YAP revealed that, while nuclear active YAP expression is confined to basal layer cells in the WT normal urothelium, the hypertrophic urothelium of Numb-KO mice shows a diffuse distribution of cells with average YAP expression level significantly higher than that detected in WT basal urothelial cells (**Extended Data Fig. 4b**). Significantly higher active YAP expression was also detected in the side-by-side comparison of both early preneoplastic/hyperplastic and not-infiltrating CIS lesions of BBN-treated Numb-KO *vs.* WT mice (**Extended Data Fig. 4c**). This increase in nuclear YAP levels was maintained in the comparison of BBN-Numb-KO *vs.* BBN-WT secondary tumors generated by retransplantation of primary BBN-induced infiltrating tumors (**Extended Data Fig. 4d**). Collectively, these findings argue that the aberrant engagement of YAP/TAZ signaling is an early driving event in the stepwise process of Numb loss-directed spontaneous or carcinogen-induced urothelial transformation. Besides representing further evidence of the association between Numb loss and YAP hyperactivation found in human BCa cell line models, these findings underscore the suitability of the paired Numb-KO *vs.* WT and BBN-Numb-KO *vs.* BBN-WT murine models to dissect the functional contribution of YAP hyperactivation to the biology of Numb-deficient BCa. To this end, we took advantage of the ability of normal and tumor mouse bladder urothelial cells to originate *ex vivo* in a reconstituted extracellular matrix (Matrigel) self-organized 3D organotypic structures (referred to hereafter as mouse bladder organoids, MBOs), amenable to molecular and functional studies^29^. This approach enabled us to investigate in a highly tractable and pathophysiologically relevant experimental setting the phenotypic alterations linked to Numb loss/YAP hyperactivation both in the normal urothelium (comparing WT *vs.* Numb-KO MBOs) and in bladder tumors (comparing BBN-WT *vs.* BBN-Numb-KO tumor MBOs).

An initial IF analysis of active YAP expression in MBOs generated from aged-matched 16-week-old Numb-KO and WT mice revealed that the absence of Numb was associated with a diffuse distribution of cells with high YAP intranuclear levels across the entire organoid population (**Fig. 3a**), mirroring the findings from the *in situ* analysis of the Numb-KO *vs.* WT bladder mucosa (**Extended Data Fig. 4b**). In addition, gross morphological differences appeared evident between the two MBO cultures. WT-MBOs showed a typical round-shaped morphology and a smooth surface deprived of invasive protrusions (**Fig. 3b**). In contrast, Numb-KO MBOs appeared as rapidly growing, irregular, multilobular structures with numerous budding protrusions infiltrating the surrounding matrix, indicative of an invasive phenotype (**Fig. 3b**). To quantify these morphological differences, we assessed well-defined physical parameters which are informative of key biological processes that determine the final organoid morphology^30–32^ (see also Materials and Methods): *i)* overall surface area – cell growth and proliferation potential, *ii*) circularity/roundness – degree of differentiation/maturation and strength of cell-cell contacts, *iii*) roughness and shape complexity – cell invasion potential. From these analyses, we concluded that the absence of Numb is associated with an aberrant morphogenetic program, characterized by hyperplasia (increased area), aberrant differentiation and loss of cell-to-cell cohesion (decreased circularity/roundness), and emergence of an invasive phenotype (increased roughness/shape complexity) (**Fig. 3c**).

**Figure 3.**
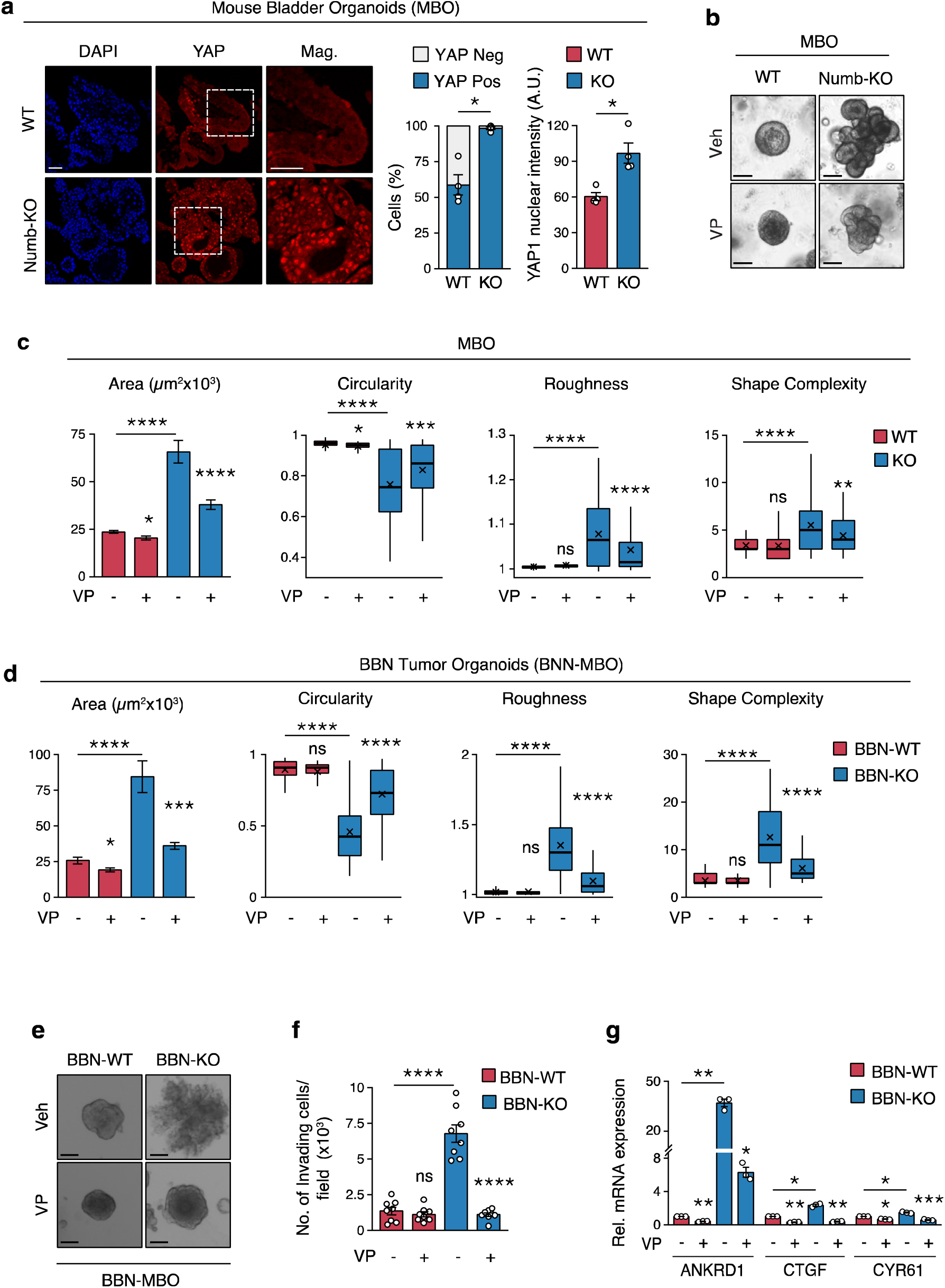
Numb loss induces invasive phenotypes and YAP signaling activation in normal and tumor urothelial cells. **a.** Left, Representative confocal IF images of endogenous YAP expression in FFPE sections of WT and Numb-KO mouse bladder organoids (MBOs) grown in 3D-Matrigel. Nuclei are stained with DAPI. Bars, 100 μm; Mag., 50 μm. The % of YAP-positive cells (middle) and the mean nuclear YAP intensity (arbitrary units, A.U.) (right) are expressed as mean ± SEM, n=4 fields/condition from one experiment, representative of two independent experiments. *, p<0.05 *vs.* WT by Welch’s t-test. **b.** Representative images of WT-and Numb-KO MBOs grown in 3D-Matrigel in the presence of vehicle (Veh) or 25 nM Verteporfin (VP) for 10 days. Bar, 100 μm. **c.** Analysis of morphometric parameters (Area, Circularity, Roughness, and Shape Complexity) of WT *vs*. Numb-KO MBOs treated as in ‘b’. Area (µm^2^) is reported as the mean ± SEM. Other parameters are reported, in this and other relevant panels in the figure, as boxplots delimited by 25th and 75th percentiles and showing the median (horizontal line) and the mean (X). The whiskers span from the smallest and largest data values within a 1.5 interquartile range. Data were obtained from three independent experiments. ****, p<0.0001; ***, p<0.001; **, p<0.01; *, p<0.05; ns, not significant relative to matching condition by FDR-adjusted pairwise Welch’s t-test. **d.** Cells dissociated from BBN-induced bladder tumors in WT and Numb-KO mice (BBN-WT and BBN-KO) were grown as MBOs (BBN-MBO) in 3D-Matrigel in the presence of 25 nM VP or vehicle for 7 days. Morphometric parameters were assessed and reported as in ‘c’. Data were obtained from three independent experiments. **e.** Representative images of BBN-WT and BBN-KO tumor MBOs grown as in ‘d’. Bar, 100 μm. **f.** Transwell Matrigel invasion assay of BBN-WT and BBN-KO cells treated with VP (100 nM for 18 h) or vehicle. Graph shows the number of invading cells/field expressed as the meanLJ±LJSEM of 8 microscope fields covering the entire migratory area, from two independent experiments. ****, p<0.0001; ns, not significant by FDR-adjusted pairwise Welch’s t-test. **g.** RT-qPCR analysis for the indicated YAP transcriptional targets in BBN-WT *vs*. BBN-KO cells treated with VP (100 nM for 12 h) or vehicle. Graphs show the relative mean fold expression ± SEM from three independent experiments. ***, p<0.001; **, p<0.01; *, p<0.05 *vs.* matching condition, by FDR-adjusted unpaired one-sample t-test (*vs.* reference sample) or two-sample Welch’s t-test (VP-*vs.* vehicle-treated BBN-KO samples).

To investigate the relevance of YAP activity to the aggressive morphological traits of Numb-KO MBOs, we employed verteporfin (VP), a clinically approved inhibitor of YAP transcriptional activity that works by preventing the YAP-TEAD interaction^33^. VP strongly inhibited the hyperplastic and invasive phenotypes of Numb-KO MBOs, resulting in smaller, more regularly shaped organoids (**Fig. 3b,c**).

Analogous results were obtained in the morphological comparison of MBOs generated by cells dissociated from BBN-WT and BBN-Numb-KO tumors (see also **Extended Data Fig. 4d,e**). The absence of Numb associated with remarkably more aggressive tumor phenotypes, reflected in a superior ability of BBN-Numb-KO MBOs to grow and locally infiltrate the surrounding matrix with cellular protrusion and migratory cells outside their exterior borders, which was reversed upon treatment with VP (**Fig. 3d,e**). Moreover, in the transwell Matrigel invasion assay, VP strongly inhibited the invasive/migratory potential of BBN-Numb-KO cells, reducing the number of invading cells to that of BBN-WT cells (**Fig. 3f**). RT-qPCR analysis of selected YAP targets genes, *Ankrd1, Ctgf* and *Cyr61*, confirmed the constitutively higher YAP transcriptional activity in BBN-Numb-KO *vs.* BBN-WT MBOs, which was reverted by VP treatment (**Fig. 3g**). These findings were recapitulated in the GSEA of the global transcriptomic profiles of BBN-Numb-KO *vs.* BBN-WT MBOs using a YAP/TAZ transcriptional target signature^16^ (**Extended Data Fig. 5a**). Remarkably, in line with results obtained in human BCa cell lines, GSEA analysis revealed a significant enrichment of an EMT signature (from Cancer “Hallmarks”) in BBN-Numb-KO *vs.* BBN-WT MBOs (**Extended Data Fig. 5a**), which was also sensitive to VP treatment (**Extended Data Fig. 5a**). This was an unexpected finding indicating that the acquisition of plasticity/EMT traits in Numb-deficient BCa cells, likely involved in their aggressive invasive phenotypes, is a downstream consequence of YAP hyperactivation.

We then asked whether genetic silencing of YAP phenocopies the inhibitory effects of VP treatment on the aggressive morphological traits of BBN-Numb-KO tumor MBOs. Stable KD of YAP using a lentiviral shRNA vector inhibited the hyperproliferative and invasive/migratory phenotype of BBN-Numb KO cells, similarly to VP, but had no effect on BBN-WT MBOs (**Extended Data Fig. 5b-e)**. Moreover, to provide definitive evidence of the dependency of the molecular and functional phenotypes of BBN-Numb-KO MBO cells on Numb loss, we restored Numb expression in these cells using a lentiviral Numb-GFP vector. Ectopic Numb expression inhibited YAP nuclear translocation, with evident YAP cytoplasmic retention (**Extended Data Fig. 6a,b**), and impaired YAP transcriptional activity (**Extended Data Fig. 6c**). These molecular changes were accompanied by a significant reversion of the aberrant morphological traits of BBN-Numb-KO MBOs, resulting in BBN-WT-like morphological structures (**Extended Data Fig. 6d**). Together, these data demonstrate that, through a molecular mechanism requiring hyperactivation of YAP signaling, loss of Numb drives an aberrant morphological program in the normal urothelium that precedes its overt malignant transformation, while conferring a higher degree of biological aggressiveness in already transformed BCa tumor cells by enhancing their proliferative and invasive/migratory potential, likely through the induction of a YAP-dependent EMT program.

The increased nuclear localization and transcriptional activity of YAP observed in cells lacking Numb, prompted us to investigate the status of Hippo signaling cascade. Core components of this pathway are the serine/threonine kinases, MST1 and LATS, whose sequential activation leads to the phosphorylation and subsequent cytoplasmic sequestration of YAP, ultimately repressing its nuclear translocation and transcriptional activity^21,34^. We therefore hypothesized that Hippo signaling could be repressed in Numb-deficient BCs. By immunoblotting of cellular lysates from BBN-Numb-KO *vs.* BBN-WT MBOs, we detected markedly reduced levels of the phosphorylated active forms of MST1 and LATS in BBN-Numb-KO cells compared with BBN-WT cells, despite having similar levels of total protein (**Extended Data Fig. 6e**). This reduction in the phosphorylated Hippo pathway kinases was in line with the observed reduction in inactive phosphorylated YAP in BBN-Numb-KO *vs.* BBN-WT cells (**Extended Data Fig. 6e**).

These results demonstrate that hyperactivation of the YAP/TAZ pathway subsequent to Numb loss is mechanistically linked to the downregulation of the upstream canonical Hippo signaling cascade.

### YAP hyperactivation downstream of Numb loss is dependent on RhoA/ROCK signaling

The Hippo-YAP signaling pathway operates as a nexus that, in response to microenvironmental cues, controls multiple cellular and context-specific responses essential for tissue homeostasis, including proliferation, differentiation, cell plasticity and stemness^34,35^. A wide range of architectural and mechanical cues, transmitted through cell-cell junctions and cell-matrix adhesions, as well as multiple extracellular ligands/growth factors and downstream signaling pathways, can control this pathway through complex canonical and non-canonical arms^36–38^. In this context, both upstream and downstream events linked to actin dynamics and cytoskeletal organization, play a pivotal role in the regulation of Hippo-YAP signaling^34,35,38,39^. This is clearly demonstrated by the ability of the F-actin disrupting agent, Lantruculin, to impede YAP/TAZ activation in the context of both canonical or non-canonical arms of Hippo-YAP signaling regulation^36,40,41^. On these bases, to gain a deeper understanding of the mechanism driving YAP hyperactivation following Numb loss, we leveraged the BBN Numb-KO *vs.* WT MBO model to perform an immunofluorescence-based phenotypic screening using pharmacological inhibitors of key regulators of actin dynamics and cytoskeleton organization. In particular, we focused on the small GTPases, RhoA and Rac1, guided by previous evidence indicating that loss of Numb control over their activity results in alterations in actin dynamics and induces different types of cell motility phenotypes^42–44^. In this analysis, we monitored YAP intranuclear translocation *vs.* cytoplasmic retention in BBN Numb-KO *vs.* BBN WT cells treated with an inhibitor of RhoA (bacterial exoenzyme C3 transferase), Rac1 (RACi, NSC23766), Rho-associated protein kinase (ROCKi, Y-27632), and actin polymerization (Latrunculin A: LatA)^36^. While Rac1 inhibition had no effect on YAP nuclear translocation in BBN Numb-KO cells, the inhibition of RhoA and its downstream effector ROCK reduced nuclear YAP levels similarly to LatA (**Fig. 4a and Extended Data Fig. 7**). In contrast, no significant effects were observed in BBN WT (**Fig. 4a and Extended Data Fig. 7**). These results point to the involvement of the RhoA/ROCK actin regulatory pathway in the control of YAP signaling by Numb.

**Figure 4.**
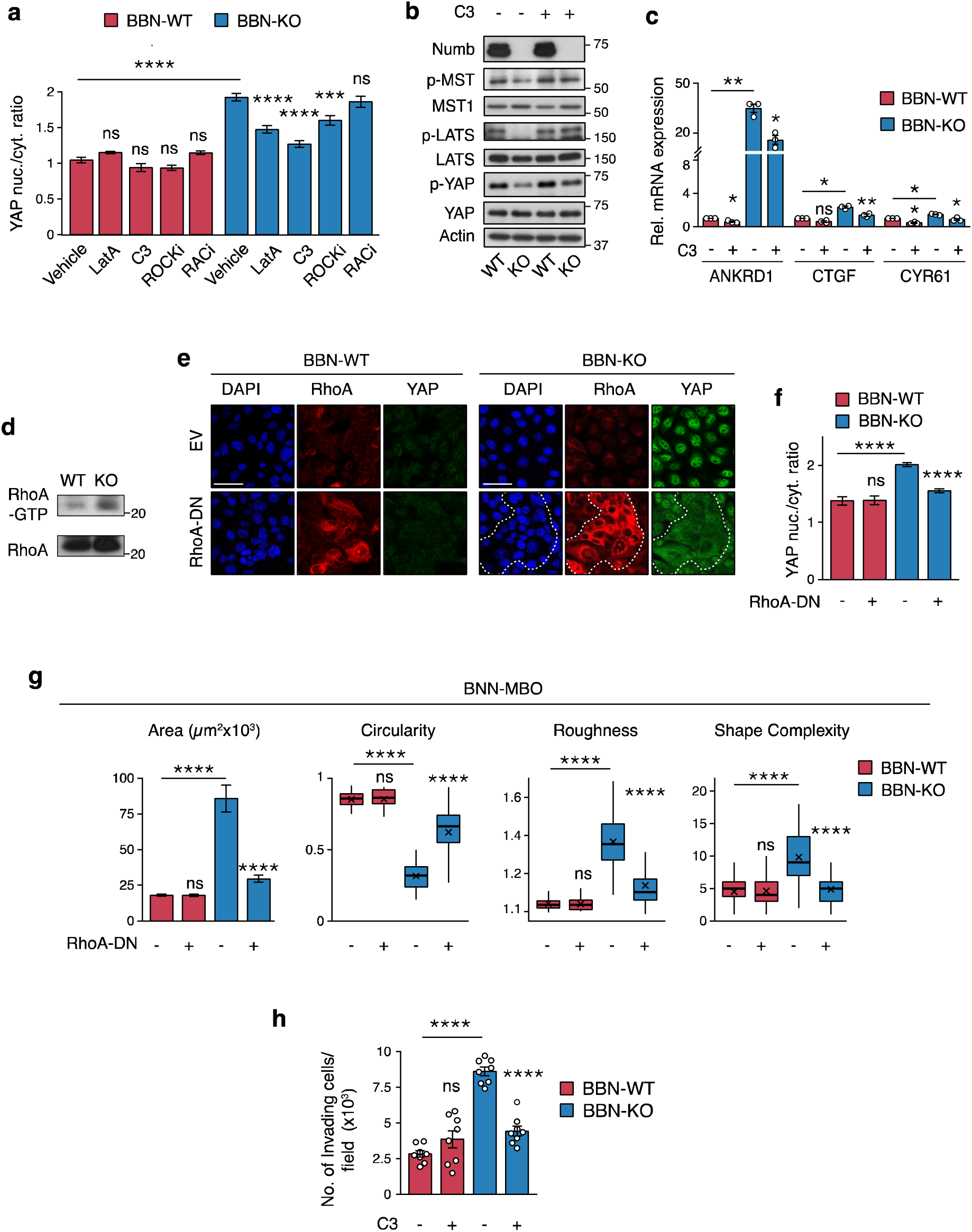
Rho-A/ROCK-dependent actin remodeling is involved in YAP hyperactivation downstream of Numb loss. **a**. BBN-WT and BBN-KO tumor cells were treated with the actin polymerization inhibitor LatA (500 nM for 6 h), Rho-A inhibitor C3 transferase (3 µg/ml for 6 h), ROCK inhibitor Y-27632 (ROCKi, 10 µM for 12 h), Rac1 inhibitor NSC-23766 (RACi, 10 µM for 12 h) or vehicle and co-stained for YAP, Numb and DAPI. Quantification of YAP nuclear/cytoplasmic ratio is shown. Graphs show the mean/field ± SEM, n=11 fields/condition, from two independent experiments. ****, p<0.0001; ***, p<0.001; ns, not significant, relative to matching controls by Tukey’s HSD test. Representative confocal images are shown in Extended Data Fig. 7. **b.** Immunoblot analysis of the expression levels and phosphorylation status of Hippo pathway components (MST, LATS, YAP) in BBN-WT *vs*. BBN-KO cells, treated with C3 transferase (3 µg/ml, 6 h) or vehicle. Numb expression was verified. Actin, loading control. Blots shown are representative of two independent experiments. **c.** RT-qPCR for the indicated YAP transcriptional targets in BBN-WT *vs*. BBN-KO cells treated as in ‘b’. Graphs show the relative mean fold expression ± SEM from three independent experiments. **, p<0.01; *, p<0.05; ns, not significant *vs.* matching condition, by FDR-adjusted unpaired one-sample t-test (*vs.* reference sample) or two-sample Welch’s t-test (C3-*vs.* vehicle-treated BBN-KO samples). **d**. Pull-down assay of activated RhoA in BBN-WT *vs*. BBN-KO cells. The amount of RhoA-GTP protein bound to beads and total RhoA in cell lysates was measured by immunoblot using an anti-RhoA antibody. Blots are representative of two independent experiments. **e.** Representative confocal fluorescence images of BBN-WT and BBN-KO tumor cells transduced with a lentiviral vector encoding dominant negative RhoA (RhoA-DN) or empty vector (EV) and co-stained for RhoA, YAP and DAPI. Bars, 50 µm. Dashed line delineates a cluster of RhoA/DN overexpressing cells in BBN-KO cells. **f.** Quantification of YAP nuclear/cytoplasmic ratio in BBN-WT-and BBN-KO cells treated as in ‘e’. Graphs show the mean/field ± SEM, n=20 fields/condition, from two independent experiments. ****, p<0.0001; ns, not significant, relative to matching controls by Tukey’s HSD test. **g.** Morphometric analysis of tumor BBN-MBOs generated from cells described in ‘e’ (see legend to Fig. 3c for the quantification of morphometric parameters). Data were obtained from three independent experiments. ****, p<0.0001; ns, not significant, relative to matching condition by FDR-adjusted pairwise Welch’s t-test. **h.** Transwell Matrigel invasion assay of BBN-WT and BBN-KO cells treated with C3 transferase (3 µg/mL, 18h) or vehicle. Graph shows the number of invading cells/field expressed as the meanLJ±LJSEM of 8 microscope fields covering the migration area, from two independent experiments. ****, p<0.0001; ns, not significant, relative to matching controls by FDR-adjusted pairwise Welch’s t-test.

Further investigation of the role of RhoA revealed that its inhibition with C3 transferase induced reactivation of the YAP-inhibitory Hippo pathway selectively in BBN Numb-KO *vs.* BBN WT cells, as evidenced by the increased phosphorylation levels of MST1, LATS and YAP (**Fig. 4b**). In addition, C3 transferase reduced the expression of YAP transcriptional targets (**Fig. 4c**). Thus, it appears that active RhoA is required for YAP hyperactivation in the Numb-KO condition. In line with this idea, we showed that BBN Numb-KO cells display higher levels of the active GTP-bound form of RhoA compared with BBN-WT cells (**Fig. 4d**). Moreover, expression of a dominant-negative RhoA mutant (TN19-RhoA) in BBN Numb-KO tumor cells inhibited YAP nuclear translocation (**Fig. 4e,f**) and reversed the phenotypes associated with biological aggressiveness, as witnessed in the morphological analysis of MBOs (**Fig. 4g**). In contrast, no effects of dominant-negative RhoA were observed in BBN-WT tumor cells (**Fig. 4e-g**). RhoA inhibition with C3 transferase also reduced the invasive/migratory potential of BBN Numb-KO tumor cells in the transwell Matrigel invasion assay, to levels similar to BBN-WT cells (**Fig. 4h**). Notably, none of the above phenotypes were affected by pharmacological (RACi, NSC-23766 inhibitor) or genetic (T17N dominant-negative mutant) inhibition of Rac1 activity (**Extended Data Fig. 8a-e**).

Next, we examined in more detail the involvement of the RhoA downstream effector, ROCK, which signals to the actin cytoskeleton and contractile machinery by regulating the phosphorylation status of two proteins: cofilin, a protein that in its phosphorylated form becomes inactive and loses its function in actin filament disassembly^45^; myosin light chain 2 (MLC2), a direct phosphorylation target of ROCK involved in the regulation of actomyosin contractility and stress fiber assembly/contraction^46,47^. We found increased levels of phosphorylated cofilin and MLC2 in BBN Numb-KO *vs.* BBN WT, indicating that loss of Numb is associated with higher basal ROCK activity (**Fig. 5a-c**). Using the ROCK inhibitor (ROCKi, Y-27632), we verified that the high levels of phosphorylated MLC2 (pMLC2) in BBN Numb-KO cells were dependent on ROCK activity (**Fig. 5b,c**).

**Figure 5.**
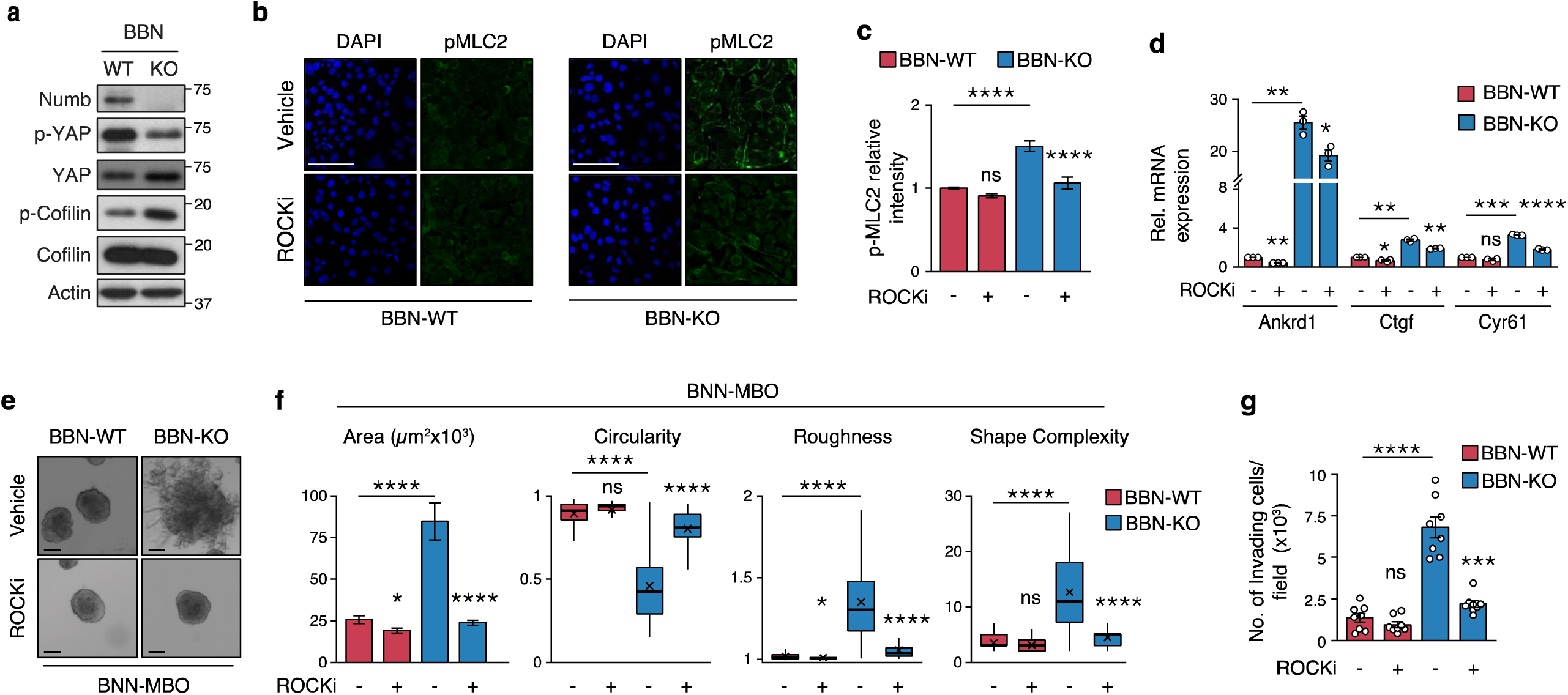
ROCK inhibition prevents YAP hyperactivation induced by loss of Numb. **a.** Immunoblot analysis of total and phosphorylated levels of cofilin and YAP in BBN-WT *vs*. BBN-KO cells. Actin, loading control. Data are representative of two independent experiments. **b.** Representative confocal images of BBN-WT and BBN-KO cells treated with vehicle or ROCK inhibitor Y-27632 (10 µM, 12 h) and co-stained for the phosphorylated form of MLC2 (pMLC2) and DAPI. Bar, 50 µm. **c.** Quantification of the experiment in ‘b’. Graphs show the relative intensity of pMLC2 in BBN-KO *vs*. BBN-WT cells expressed as the mean/field ± SEM, n=18 random fields/condition from two independent experiments. ****, p<0.0001; ns, not significant, relative to matching controls by Tukey’s HSD test. **d.** RT-qPCR of the indicated YAP transcriptional targets in BBN-KO *vs.* BBN-WT cells treated with ROCKi Y-27632 (10 µM, 8 h) or vehicle. Graphs show the relative mean fold expression ± SEM from three independent experiments. ****, p<0.0001; ***, p<0.001; **, p<0.01; *, p<0.05; ns, not significant, *vs.* matching condition by FDR-adjusted unpaired one-sample t-test (*vs.* reference sample) or two-sample Welch’s t-test (ROCKi *vs.* vehicle BBN-KO samples). **e.** Representative images of tumor BBN-MBOs generated from BBN-WT and BBN-KO cells treated with ROCKi (Y-27632, 10 µM). Bars, 100 µm. **f.** Morphometric analysis of the indicated parameters in BBN-MBOs derived from cells described in ‘e’ (see also legend to Fig. 3c). Data were obtained from three independent experiments. ****, p<0.0001; *, p<0.05; ns, not significant, relative to matching condition by FDR-adjusted pairwise Welch’s t-test. **g.** Transwell Matrigel invasion assay of BBN-WT and BBN-KO cells treated with ROCKi Y-27632 (10 µM, 18 h) or vehicle. Graph shows the number of invading cells/field expressed as the meanLJ±LJSEM of 8 random microscope fields covering the entire migration area, from two independent experiments. ****, p<0.0001; ns, not significant, relative to matching controls by FDR-adjusted pairwise Welch’s t-test.

Finally, we demonstrated that ROCK has a key role in YAP signaling regulation downstream of Numb, as evidenced by the reduction of YAP transcriptional activity in BBN Numb-KO MBOs treated with Y-27632 (**Fig. 5d**). In addition, ROCK inhibition reversed the aggressive phenotypes of BBN-Numb-KO tumor cells, as witnessed in the analysis of morphological parameters measuring MBO growth and invasive/migratory potential (**Fig. 5e-g**) .

Together, these results point to the selective dependency of the aggressive invasive/migratory phenotype of BBN Numb-KO tumor cells on the aberrant activation of a RhoA/ROCK axis signaling to the actin cytoskeleton. Therapeutic targeting of this circuitry could therefore serve as a strategy to treat aggressive Numb-deficient BCa.

### Dysregulation of RhoA/ROCK/YAP signaling underlies the invasive phenotype of Numb-deficient human BCa cells

To investigate the relevance of RhoA/ROCK/YAP signaling to Numb-deficient human BCa, we initially employed the human RT4 cell line, which represents a model of low aggressive, well-differentiated non-invasive luminal BCa^48–50^. The comparison of Numb-KD *vs.* CTR-KD RT4 BCa cell lines confirmed the results obtained in the BBN Numb-KO *vs.* BBN WT mouse model, showing that the absence of Numb leads to YAP hyperactivation through the downregulation of the canonical Hippo pathway, as witnessed by the reduced phosphorylation levels of MST1, LATS, and YAP in Numb-KD *vs.* CTR-KD RT4 cells (**Fig. 6a**). Moreover, several lines of evidence support the finding that activation of RhoA/ROCK signaling is upstream of YAP hyperactivation in Numb-deficient RT4 cells. Indeed, pharmacological inhibition of RhoA and ROCK, but not Rac1, reduced the nuclear/cytoplasmic YAP ratio selectively in Numb-KD RT4 cells, mimicking the effect of the actin polymerization inhibitor LatA (**Fig. 6b and Extended Data Fig. 9a**). The increased cytoplasmic retention of YAP correlated with increased levels of inactive phosphorylated YAP, seemingly due to restoration of the Hippo pathway activity, indicated by increased p-MST1/2 levels, thus mirroring the results observed in Numb-KD RT4 cells treated with the RhoA inhibitor C3 transferase (**Fig. 6c**). In addition, Numb-KD *vs.* CTR-KD RT4 cells displayed higher constitutive levels of activated ROCK, as witnessed by the higher levels of the phosphorylated forms of its downstream effectors, cofilin and MLC2 (**Fig. 6d and Extended Data Fig. 9b,c**). Treatment with RhoA and ROCK inhibitors (C3 transferase and Y-27632, respectively) also strongly inhibited the transcription of the YAP target genes, *ANKRD1, CTGF* and *CYR6* (**Fig. 6e**), similarly to the YAP transcriptional inhibitor VP (**Extended Data Fig. 9d**). Therefore, it appears that the same molecular circuitry involving Rho/ROCK, actin remodeling and Hippo signaling is required for the regulation of YAP by Numb in both mouse and human BCa cells.

**Figure 6.**
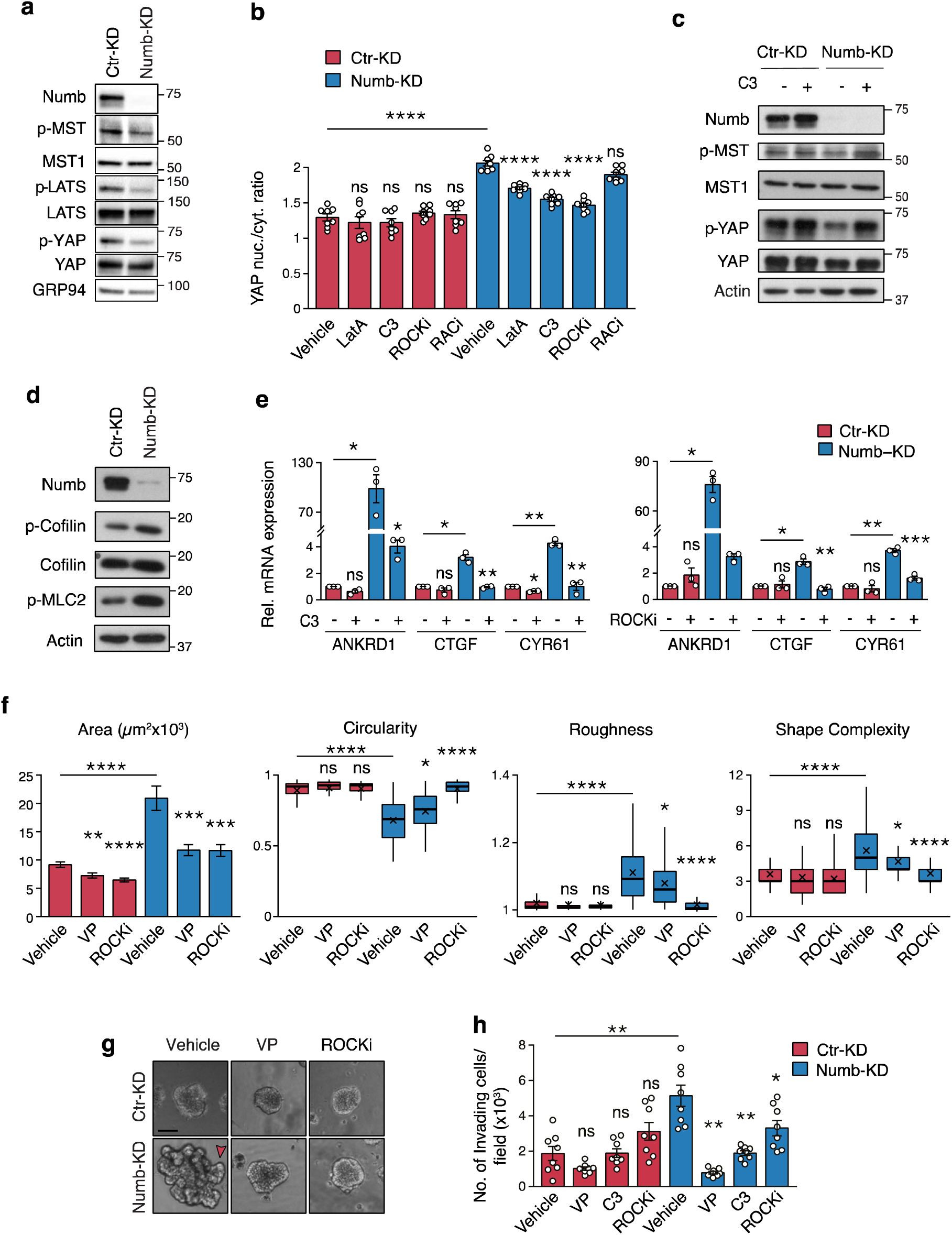
Loss of Numb triggers RhoA/ROCK-dependent YAP hyperactivation in human RT4 BC cells. **a.** Immunoblot analysis of Numb and Hippo pathway components (MST, LATS, YAP) in control-(Ctr-KD) and Numb-silenced (Numb-KD) RT4 cells. Grp94, loading control. Data are representative of two independent experiments. **b.** Ctr-KD and Numb-KD RT4 cells were treated with actin polymerization inhibitor Latrunculin-A (LatA, 500 nM for 6 h), Rho-A inhibitor C3 transferase (C3, 3 µg/ml for 6 h), ROCK inhibitor Y-27632 (ROCKi, 10 µM for 12 h), Rac1 inhibitor NSC-23766 (RACi, 10 µM for 12 h) or vehicle and co-stained for endogenous YAP and DAPI. Representative confocal images are shown in Extended Data Fig. 9a. Quantification of YAP nuclear/cytoplasmic ratio is shown as the mean/field ± SEM, n=8 fields/condition, from two independent experiments. ****, p<0.0001; ns, not significant, relative to matching controls by Tukey’s HSD test. **c.** Immunoblot analysis of Numb, and total and phosphorylated YAP and MST1 in Ctr-KD *vs.* Numb-KD RT4 cells treated with vehicle or C3 transferase (3 μg/mL, 6 h). Actin, loading control. Data are representative of two independent experiments. **d.** Immunoblot analysis of Numb, total and phosphorylated cofilin, and phosphorylated MLC2 (p-MLC2), in Ctr-KD *vs.* Numb-KD RT4 cells. Actin, loading control. Blots are representative of two independent experiments. **e.** RT-qPCR of the indicated YAP transcriptional targets in Numb-KD *vs.* Ctr-KD RT4 cells treated with C3 (3 µg/ml, 6 h) (left) or ROCKi Y-27632 (50 µM, 8 h) (right), or vehicle. Graphs show the relative mean fold expression ± SEM from three independent experiments. ***, p<0.001; **, p<0.01; *, p<0.05; ns, not significant, *vs.* matching condition by FDR-adjusted unpaired one-sample t-test (*vs.* reference sample) or two-sample Welch’s t-test (drug *vs.* vehicle Numb-KD samples). **f.** Analysis of morphometric parameters in 3D-Matrigel organoids derived from Ctr-KD and Numb-KD RT4 cells treated with vehicle (Veh), Verteporfin (VP, 25 nM) or ROCKi Y-27632 (10 µM). Area (μm^2^) is reported as mean ± SEM. Other parameters are reported as boxplots delimited by 25^th^ and 75^th^ percentiles and showing the median (horizontal line) and the mean (X). The whiskers span from the smallest and largest data values within a 1.5 interquartile range. Data were obtained from two independent experiments. ****, p<0.0001; ***, p<0.001; *, p<0.05; ns, not significant, relative to matching condition by FDR-adjusted pairwise Welch’s t-test. **g.** Representative images of organoids derived from Ctr-KD *vs.* Numb-KD RT4 cells treated as in ‘f’. Red arrows point to invasive protrusions in Numb-KD RT4 cells. Bar, 100 µm. **h.** Transwell Matrigel invasion assay of Ctr-KD and Numb-KD RT4 cells treated with vehicle, VP (100 nM), C3 (3 µg/mL) and ROCKi Y-27632 (10 µM) for 48 h. Graph shows the number of invading cells/field expressed as the meanLJ±LJSEM of 8 microscope fields from two independent experiments. **, p<0.01; *, p<0.05; ns, not significant vs. matching condition by FDR-adjusted pairwise Welch’s t-test.

Next, we verified whether this mechanism could be relevant to biological phenotypes induced by Numb loss in human BCa, analyzing the effects of the different pathway inhibitors on the morphology and invasion/migration potential of 3D-Matrigel organoids generated by Numb-KD *vs.* CTR-KD RT4 cells. Similarly to the BBN tumor model, we found that the absence of Numb exacerbates the biological aggressiveness of RT4 cells, as evidenced in the comparison of Numb-KD *vs.* CTR-KD RT4 organoids by the increase in organoid size (area), propensity to form invading protrusions (**Fig. 6f,g**), and increased invasive/migratory potential in the transwell Matrigel invasion assay (**Fig. 6h**). All these phenotypes were reversed selectively in Numb-KD RT4 cells by pharmacological inhibition of YAP (VP) and RhoA/ROCK (C3/Y-27632) signaling (**Fig. 6f-h**).

This set of molecular and functional findings obtained in the RT4 BCa cell line was entirely recapitulated following Numb silencing in another model of low aggressive, moderately differentiated non-invasive human BCa, the RT112 cell line^48–50^ (**Extended Data Fig. 10a-g**). Together, these data, obtained in two independent human NMIBC cell line models with proficient Numb expression and low degree of aggressiveness, further support the notion that loss of Numb results in the acquisition of biological phenotypes associated with aggressive BCa, establishing its relevance to the human BCa disease. Mechanistically, these data are further evidence that the aggressive biology of Numb-defcient BCa cells depends on the aberrant activation of RhoA/ROCK signaling to the actin cytoskeleton, downstream of Numb loss, which leads to suppression of the YAP-inhibitory Hippo pathway and consequently YAP hyperactivation. Remarkably, this molecular circuitry, functionally validated in the above human NMIBC cell lines, appears to be relevant to real-life NMIBC patients, since we detected significantly higher levels of phosphorylated cofilin, indicative of increased ROCK activity, in Numb^Low^ NMIBC TUR samples, compared with Numb^High^ (**Fig. 7a,b**). This finding is also in keeping with the increased nuclear accumulation and transcriptional activity of YAP that we observed in NMIBC patients with deficient Numb status (see **Fig. 1h-j**).

**Figure 7.**
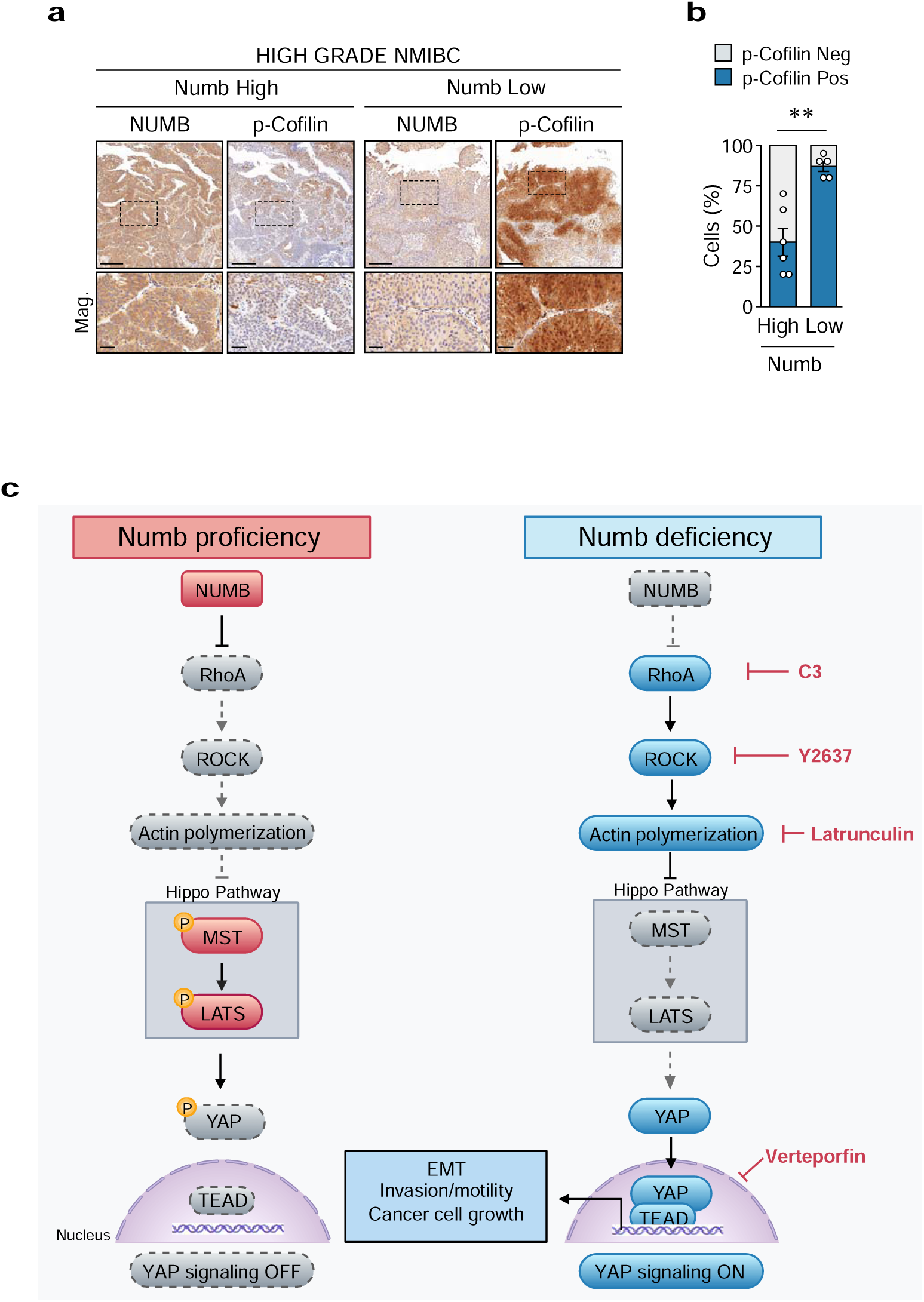
Loss of Numb expression correlates with increased p-cofilin levels in NMIBC patients. a. Representative IHC staining of phosphorylated cofilin (p-Cofilin) and Numb in Numb^High^ and Numb^Low^ high-grade NMIBC TUR specimens. Magnifications (Mag.) of the boxed areas are shown in the lower panels. Bars, 500 µm; Mag, 100 µm. **b.** Quantification of the % of p-cofilin positive cells in 6 Numb^High^ *vs.* 5 Numb^Low^ high-grade NMIBC TUR specimens. **, p=0.0019 by Welch’s *t*-test. **c.** Schematic representation of the molecular events influencing YAP activation state in the bladder urothelium in Numb proficient or deficient conditions. In Numb proficient conditions (Numb proficiency), the presence of Numb keeps in check RhoA/ROCK signaling to the actin machinery, leading to activation of the Hippo pathway, which phosphorylates YAP resulting in its cytoplasmic retention and inactivation (YAP signaling OFF). In Numb deficient conditions (Numb deficiency), the absence of Numb leads to activation of RhoA/ROCK signaling to the actin machinery, which in turn suppresses the Hippo pathway, resulting in nuclear translocation of unphosphorylated YAP and transcription of its target genes via interaction with TEAD (YAP signaling ON). These transcriptional changes, likely through the induction of an EMT program, promote the acquisition of proliferative and invasive/migratory phenotypes underlying the biological aggressiveness of Numb-deficient bladder tumors.

Together, these findings point to the RhoA/ROCK/YAP signaling pathway as a potential actionable vulnerability for therapeutic intervention in Numb-deficient NIMBCs.

## DISCUSSION

In this study, we have uncovered that loss of the tumor suppressor Numb is a hallmark of aggressive disease course in real-life BCa patients and identified key mechanisms causal to the biological aggressiveness of Numb-deficient bladder tumorigenesis, highlighting the potential of Numb loss as a prognostic and therapeutic biomarker for personalized, risk-adjusted treatment of BCa patients, in particular NMIBC patients.

Numb is an evolutionarily conserved cell fate determinant and endocytic adaptor protein^51^ that has emerged in recent years as a tumor suppressor protein with prognostic and pathogenetic relevance in various cancers, such as breast, prostate, brain, lung, and colon cancer^17–19,22,44,52–56^. The tumor suppressor function of Numb has been extensively characterized in breast cancer, where its loss of function leads to the emergence and uncontrolled expansion of cancer stem cells through inhibition of the tumor suppressor p53 and increased activity of the oncogene Notch^17–19,22,52,53^.

In this study, using a range of experimental systems, including established human BCa cell lines, *in vivo* mouse models, primary organoid cultures and patient cohorts, we have established that Numb is a clinically relevant tumor suppressor protein in the bladder. Retrospective studies of MIBC and NMIBC patient cohorts demonstrated that low Numb expression in the tumor predicts worse overall survival and increased risk of progression to MIBC, respectively. Moreover, using genetically engineered mouse models, we uncovered that Numb loss is causal in bladder tumorigenesis, confirming its role as a potent tumor suppressor. Indeed, transgenic deletion of the *Numb* gene in the CK5-expressing basal layer of the urothelium was alone sufficient to induce the formation of preneoplastic lesions, non-invasive tumors and muscle-invasive tumors. This finding aligns with the emerging view that intrinsically aggressive BCa with a propensity for muscle invasion originate from basal, rather than suprabasal, cells^24–26,57^. The development of a genetically engineered mouse model recapitulating the non-invasive to invasive bladder cancer transition holds particular significance in the field of bladder cancer, considering the relative paucity of *in vivo* models amenable to study the underlying biology of NMIBC to MIBC progression, which still remains largely unexplored^58^. Related to this point, the progressive range of lesions observed in our Numb-KO mouse model supports the idea that NMIBC and MIBC represent different stages along a disease continuum. This finding is in keeping with recent experimental studies in the mouse^26^ and with evidence of a high degree of similarity between genetic alterations in NMIBC and MIBC patients^14^, which have challenged the historical view of NMIBC and MIBC as *ab initio* distinct pathobiological entities^15^.

In addition to Numb loss-of-function being alone sufficient to drive spontaneous tumorigenesis, it can also cooperate with other oncogenic insults to accelerate tumor progression and fatal outcome, as clearly demonstrated in Numb-KO mice exposed to the chemical carcinogen, BBN. This compound induces bladder tumorigenesis in mice, mirroring the progression of naturally occurring human BCa disease^25,27,28^. By comparing the effects of BBN on WT and Numb-KO mice, we could formally prove that the absence of Numb exacerbates the biological aggressiveness of BCa cells by conferring a highly proliferative and invasive/migratory phenotype, as evidenced in the morphological comparison of BBN-Numb-KO *vs.* BBN-WT tumor organoids and the transwell invasion/migratory assay. These findings were entirely replicated in preclinical models of superficial non-invasive human BCa, the RT4 and RT112 cell lines^48–50^, which switched to an overtly invasive phenotype following silencing of Numb expression. Together, these results point to Numb loss as a molecular hallmark of biological aggressiveness in human BCa, strongly supporting our clinical evidence of Numb loss as a biomarker of clinically aggressive disease course in real-life BCa patients.

At the mechanistic level, we demonstrated that the aggressive biological phenotypes conferred by Numb loss are dependent on a RhoA/ROCK/Hippo molecular circuitry leading to hyperactivation of YAP (**Fig. 7c**). In the presence of functional Numb, RhoA/ROCK signaling to the actin cytoskeleton is restrained, allowing an active Hippo pathway to inhibit the activity of YAP through its phosphorylation and cytoplasmic retention (**Fig. 7c**). In contrast, in the absence of Numb, RhoA/ROCK signaling is upregulated, leading to suppression of the Hippo pathway via actin cytoskeleton remodeling and consequently nuclear translocation of YAP and activation of YAP/TAZ transcriptional activity (**Fig. 7c**). These findings are in line with a recent report showing that Numb-mediated inhibition of RhoA/ROCK activity is implicated in the regulation of migration and proliferation in colon cancer cells^44^. Therefore, the dysfunction of the Numb-RhoA/ROCK/YAP axis identified in this study may have far-reaching implications in cancer biology beyond BCa.

One open question from our study is how Numb regulates RhoA/ROCK signaling. We previously showed that the endocytic/sorting function of Numb is involved in the regulation of another small GTPase, Rac1. Numb loss results in Rac1 activation, promoting the formation of specialized actin-based lamellipodia protrusions and cell motility phenotypes^42,43^. These events are associated with the relocalization of Rac1 to the plasma membrane through an EFA6B-ARF6-dependent endocytic recycling route, which is negatively regulated by the direct interaction of Numb with the guanine nucleotide exchange factor (GEF) EFA6B^43^. Therefore, Numb might control RhoA subcellular localization and activation state by interacting with positive (GEFs) or negative (GAPs: GTPase-activating proteins; GDIs: guanine nucleotide dissociation inhibitors) regulators of RhoA activity. This hypothesis is in keeping with previous reports linking RhoA activation to its increased plasma membrane localization and decreased cytosolic distribution^59,60^.

Furthermore, our results showing that a RhoA-dependent/Rac1-independent mechanism is involved in Hippo pathway inactivation/YAP activation in Numb-deficient BCa, point to actin cytoskeleton remodeling events selectively controlled by RhoA/ROCK, such as actomyosin contractility and stress fiber formation^45,61^. This hypothesis, supported by our evidence of increased phosphorylation of MLC2 and cofilin in Numb-deficient BCa cells, is consistent with reports that RhoA-induced stress fibers can suppress the Hippo pathway and activate YAP^62,63^. It has been proposed that stress fibers likely act as a scaffold for several Hippo pathway components, such as AMOT and MST1/2^64–67^, and that MST1/2 is activated upon F-actin depolymerization^67^.

On the other hand, Numb has been reported to directly or indirectly interact with other actin cytoskeleton regulators, such as α-catenin, β-catenin, NF2/Merlin, AMOT and SPTAN1, which can influence the Hippo pathway’s MST1/2-LATS kinase cascade to inhibit YAP/TAZ activity^66,68,69^. Whether such mechanisms are at play in Numb-deficient BCa and how they are integrated with the Numb/RhoA/ROCK axis remain to be elucidated. However, it is apparent that Numb loss has the potential to influence multiple networks involved in actin cytoskeleton remodeling and cell geometry regulation, including tight junctions, adherens junctions and cell polarity complexes, as well as extracellular diffusible signals^34,36–40^, which could all contribute to the aggressive invasive phenotypes through RhoA/ROCK signaling activation.

The results of this study have potential clinical implications. The functional characterization of a RhoA/ROCK/YAP circuitry responsible for the aggressive biology of Numb-deficient BCa opens up opportunities for the development of targeted therapies, in particular for the treatment of NMIBC patients at high risk of MIBC progression. Indeed, our work, involving the use of pharmacological inhibitors or genetic inhibition, highlights the RhoA/ROCK/YAP axis as an actionable vulnerability which can be exploited to restrain the aggressiveness of Numb-deficient BCa (**Fig. 7c**). Notably, several drugs that can interfere with this circuitry are currently under investigation as potential anti-cancer treatments. Some of these drugs are already in clinical use, such as the YAP transcriptional activity inhibitor, VP^33^, which is employed in a broad range of ophthalmology conditions^70,71^, or the ROCK inhibitor, Fasudil, currently in clinical trials for different types of vascular and neurodegenerative disorders^72^. However, predictive biomarkers are still needed to expand their clinical indications and minimize their systemic toxicity. In the case of NMIBC, the translation of Numb status into the clinical practice would not only provide a novel biomarker of aggressive disease course; but also allow the identification of patients who might benefit from RhoA/ROCK/YAP targeted treatments.

## MATERIALS AND METHODS

### Materials

Antibodies were obtained from the following sources: anti-Numb mouse monoclonal antibody (clone Ab21, 1:1000 for IB) (#MABN2311 from Sigma Aldrich); anti-Numb rabbit monoclonal (clone C29G11) (#2756 from Cell Signaling Technologies, 1:500 for IF, 1:150 for IHC); anti-phospho MST1(Thr183)/MST2(Thr180) (#49332 from Cell Signaling Technology, 1:1000 for IB); anti-MST1 (#3682 from Cell Signaling Technology, 1:1000 for IB); anti-phospho LATS1 (Thr1079) (#PA5-105442 from Invitrogen, 1:1000 for IB); anti-LATS1 (#3477 from Cell Signaling Technology, 1:1000 for IB); anti-phospho YAP (Ser127) (#4911 from Cell Signaling Technology, 1:1000 for IB); anti-YAP1 (63.7) (#sc-101199 from Santa Cruz, 1:1000 for IB, 1:200 for IF); anti-YAP (D8H1X) (#14074 from Cell Signaling Technology, 1:200 for IF); anti-active YAP (EPR19812) (#ab205270 from Abcam, 1:500 for IF); anti-phospho Myosin Light Chain 2 (Ser19) (pMLC2) (#3671 from Cell Signaling Technology, 1:1000 for IB; 1:200 for IF); anti-phospho Cofilin-1 (Ser3) (F-11) (#sc-365882 from Santa Cruz, 1:1000 for IB, 1:300 for IF); anti-Cofilin (D3F9) (#5175 from Cell Signaling Technology, 1:1000 for IB); anti-RAC1 clone 102/Rac1 (RUO) from BD Biosciences, 1:350 for IF; anti-RhoA (26C4) (#sc-418 from Santa Cruz, 1:400 for IF); anti-RhoA (#ARH03 from Cytoskeleton, 1:2000 for IB); anti-actin (A4700 from Merck Life Sciences, 1:10000 for IB); anti-Vinculin (#V9131 from Merck Life Sciences, 1:10000 for IB); anti-CK5 (EP1601Y) (ab52635 from Abcam, 1:2000 for mice IHC, 1:200 for human IHC); anti-CK14 (LL002) (ab7800 from Abcam, 1:1000 for IF); anti-CK7 (RCK105) (ab9021 from Abcam, 1:500 for IF); anti-CK20 (ab118574 from Abcam, 1:200 for IF and IHC); Bond Polymer Refine Detection Kit (#DC9800; Leica Biosystems); Bond Primary Ab Diluent (#AR9352; Leica Biosystems); goat anti-rabbit IgG (H+L)-HRP conjugate (#1706515 Bio-Rad, 1:2500 for IB); goat anti-mouse (H+L)-HRP conjugate (#1706516 Bio-Rad, 1:2500 for IB); Alexa Cy3-and 488-conjugated secondary antibodies from Jackson ImmunoResearch (#715-165-150 and #715-545-152, respectively; 1:500 for IF); Alexa Fluor 555 donkey anti-mouse IgG (1:400 for IF) and Alexa Fluor 488 donkey anti-rabbit IgG (1:500 for IF) secondary antibodies were obtained from ThermoFisher; Mowiol-Dabco solution mounting medium (#81381 and #D27802 from Sigma-Aldrich Merck); DAPI (#32670 from Sigma-Aldrich Merck); Dispase was from Stemcell Technologies; Advanced-DMEM/F-12 was from ThermoFisher; McCoy’s 5A, RPMI and MEM mediums were from EuroClone; FGF10 (#100-26) and FGF7 (#100-19) from Peprotech; A83-01 from Sigma-Aldrich Merck; B27 from Invitrogen; Tryple Express from Gibco; Matrigel growth factor–reduced basement membrane matrix (#356231) from Corning; fetal bovine serum (FBS) from Euroclone; ROCK inhibitor Y-27632 was purchased from Selleckchem; Cell Permeable Rho Inhibitor (C3 Transferase) from Cytoskeleton Tebu-Bio; Rac1 inhibitor (NSC23766), Cell-permeable marine toxin Latrunculin A (#L5163, from Merck Life Science Srl; Verteporfin (#HY-B0146, from MedChemExpress and T3112, from TargetMol). *N*-butyl-*N*-(4-hydroxybutyl) nitrosamine (BBN) (#B8061, from Merck Life Science Srl).

### Genetically engineered mouse models

Mice were housed in pathogen-free certified animal facilities at the Cogentech Mouse Facility located at IFOM (The AIRC Institute of Molecular Oncology, Milan, Italy). All mice have been maintained in a controlled environment, at 18–23 °C, 40–60% humidity, and with 12-h dark/12-h light cycles. All animal studies were conducted with the approval of Italian Minister of Health (AUT. N. 125/2019-PR) and were performed in accordance with the Italian law (D.lgs. 26/2014), which enforces Dir. 2010/63/EU (Directive 2010/63/EU of the European Parliament and of the Council of 22 September 2010 on the protection of animals used for scientific purposes) and EU 86/609 directive and under the control of the institutional organism for animal welfare and ethical approach to animals in experimental procedures (Cogentech OPBA). The Numb-KO mouse model was generated as previously described^22^. Briefly, mice were generated by crossing Numb^lox/lox^ mice^73,74^ with CK5-Cre mice^75^. Targeted deletion of the *Numb* allele was confirmed by PCR genotyping, as described previously^73,75^.

Expression of the *Cre*-*Recombinase* allele was confirmed by PCR genotyping, using the following forward and reverse primers: CRE-FW: AACATGCTTCATCGTCGG; CRE-REV: TTCGGATCATCAGCTACACC. Male animals were used for the experiments.

### BBN-induced mouse model of bladder cancer and mouse pathology

Aged-matched 8-to 16-week-old mice were given tap water containing 0.05% BBN for 16 or 20 weeks and afterward switched to regular drinking water until the end of the experiment. The control group received tap water throughout the entire experiment. Mice treated for 16 weeks were sacrificed 2 weeks after the end of the BBN treatment (**Fig. 2c** and **Extended Data Fig. 3d**). Mice treated for 20 weeks were sacrificed upon reaching any of the predefined endpoints for high disease burden, including abdominal distension that impeded movement, loss of >15% of body weight, labored breathing, abnormal posture, hematuria (**Fig. 2d** and **Extended Data Fig. 3d**). At sacrifice, bladders were removed and stored in 4% paraformaldehyde for subsequent paraffin embedding and histological examination. The group assignment of bladder tissue sections was blinded to a board-certified pathologist for the objective histological evaluation and staging of BBN-induced lesions in the mouse urothelium. Carcinoma *in situ* (CIS) was defined as a flat tumor lesion confined to the superficial urothelial layer, with malignant urothelial cells displaying loss of cell polarity, cellular atypia, increased number of mitotic figures, and large irregular nuclei with a high nuclear/cytoplasmic ratio, according to standard pathology criteria.

### Post-cystectomy MIBC patient cohort

Archival FFPE tissue specimens were collected from a retrospective consecutive cohort of 383 patients who underwent cystectomy for bladder cancer at the European Institute of Oncology (IEO, Milan) between 1999 and 2016, following approval by the Institutional Ethical Board. The clinicopathological characteristics of this cohort are reported in **Extended Data Fig. 1a**. Numb IHC staining was performed on whole FFPE sections from 356 tumor blocks with sufficient material.

Staining intensity was scored on a scale from 0 to 3 (0, undetectable expression; 1, intermediate expression; 2-3, expression comparable to normal urothelial basal cells). The percentage of Numb-positive cells was estimated in increments of 5% over the entire tumor area within the tissue section. Tumors were classified as Numb-low if a tumor area larger than 70% exhibited an intensity score <1; otherwise, they were categorized as Numb-high.

Survival analysis was conducted using univariable and multivariable Cox proportional hazard models, with multivariable analysis adjusted for clinically relevant variables (age, gender, pT, pN, and vascular invasion). Of the 338 patients with available follow-up data, eighty patients with a pathological tumor stage of pT4 were excluded from the survival analysis to minimize the confounding effects linked to their generally very poor 5-year survival rate^76^.

### NMIBC patient cohorts

Archival FFPE transurethral resection (TUR) specimens were collected from a longitudinal series of patients diagnosed with NMIBC at the European Institute of Oncology (Milan) and Policlinico Ospedali Riuniti di Foggia (Foggia, Italy), following standard operating procedures approved by the respective Institutional Ethical Boards. Patients with the first available high-grade NMIBC lesion were selected for the analyses, excluding those patients with a synchronous muscle-invasive disease. The primary outcome was time-to-progression to MIBC in a 4-month minimum follow-up period by TUR. Patients who did not progress were right-censored at the last recorded NMIBC lesion, cystectomy, or negative cystoscopy. Complete clinicopathological information of this NMIBC cohort, including age, gender, and tumor stage, are described in detail in **Extended Data Fig. 1c**. Whole FFPE TUR sections from a total of 77 patients were IHC stained for Numb. Assessment of Numb status and univariable and multivariable analysis were performed as described above for the post-cystectomy MIBC cohort.

### Cells and culture procedures

RT4 and CLS439 cells were cultured in McCoy’s 5A medium supplemented with 10% heat-inactivated fetal bovine serum (FBS), 2mM L-Glutamine and 1% penicillin/streptomycin (P/S). RT112, KK47 and 5637 cells were cultured in RPMI medium supplemented with 10% heat-inactivated FBS, 2mM L-Glutamine and 1% P/S. HT1376 cells were cultured in MEM medium supplemented with 10% heat-inactivated FBS, 2mM L-Glutamine, 1% P/S and 0.1mM of Non-Essential Amino Acids (NEAA). Primary murine cells were grown in advanced-DMEM/F-12 supplemented with 100 ng/mL FGF10, 25 ng/mL FGF7, 500 nM A83-01, and 2% B27. BBN-induced tumor cells were maintained in 1:1 mixture of DMEM and Ham’s F12 medium, supplemented with 2 mM L-Glutamine, 5 μg/ml insulin, 0.5 μg/ml hydrocortisone, 2% B27, 20 ng/ml EGF and bFGF, and 4 μg/ml heparin. To establish mouse bladder organoids (MBOs), mouse bladder tissues were mechanically and enzymatically dissociated with 5 U/mL Dispase for 30 minutes in 5% CO_2_ humidified atmosphere at 37°C and the resulting cell suspension was dissociated to single cells with Tryple Express. Cells were then plated at a density of 2000 cells/mL for primary normal murine cells and 1000 cells/mL for BBN-induced tumor cells and human bladder cancer cell lines, and allowed to generate 3D-organoids in a drop of 50 µL of pure Matrigel in a 24-well plate for 7-10 days after seeding in a total volume of 1 mL of culture medium. Organoid cultures were incubated at 37°C in a humidified 5% CO_2_ atmosphere, with fresh medium changed every 7 days. Drug treatments started 3 days after seeding. The resulting 3D-organotypic structures were observed and photographed using an inverted phase-contrast microscope equipped with a digital camera.

### Expression, hairpin constructs and engineering of vectors

Numb knockdown in human bladder cancer cells was performed either by small interfering RNA (siRNA) or by lentiviral-driven expression of small hairpin RNA (shRNA). The efficiency of Numb silencing was assessed by Western Blotting analysis. For siRNA experiments, cells were transfected with 5 nM of control siRNA (siCTRL) or Numb siRNA (siNUMB) using Lipofectamine RNAiMAX transfection reagent (Thermo Fisher Scientific) according to the manufacturer’s instructions. A second round of transfection was performed 48 hours after the first transfection, and cells were processed for the appropriate assay 48 hours after the second transfection. The targeted sequences used were as follows: Numb siRNA, CAGCCACUGAACAAGCAGA; scrambled siRNA, AGACGAACAAGUCACCGAC^17^. pLKO.1-Numb shRNA were purchased from GE Dharmacon (oligo identification: TRCN0000007226); a hairpin against Luciferase was used as control (TRCN0000072243). pLKO.1-Yap shRNA was obtained from Addgene (#166486) and used to knockdown YAP expression in BBN-induced cancer cells. To express ectopic Numb, a pLVX-Numb-GFP construct was engineered by subcloning the Numb-GFP fragment from a pEGFP-N1/Numb construct (already available in the lab), into the EcoRI-SmaI sites of the pLVX-Puro vector (Clontech).

DNA encoding full length hRac1-DN (Rac1-T17N) was amplified by PCR from pRK5-myc/Rac1-N17 (#15904, Addgene) using the following primers: RAC1_T17N FW: TTTTTCTCGAGACGCGTATCGATTGAATTGGCCACC; RAC1_T17N RV: CTGCAGGAATTCTTACAACAGCAGGC; DNA encoding full-length hRhoA-DN (RhoA-T19N) was amplified by PCR from pSLIK/RhoA-T19N (# 84646, Addgene) using the following primers: RhoA_T19N, FW: TTTTTCTCGAGACGCGTATGGAGCAGAAGCTGATCTCCGAGG; RhoA_T19N, RV: GGATATCTGCAGAATTCACCGC. The resulting DNA fragments were inserted into the XhoI-EcoRI sites of pLVX-Puro. All constructs were sequence-verified. HEK293T were used for viral supernatants production and infection.

### Quantitative real-time PCR

Total RNA was extracted with QIAzol™ Reagent and then purified using the RNeasy kit (Qiagen). RNA reverse transcription (1μg) was performed using the High-Capacity cDNA Reverse Transcription Kit (Thermo Fisher) or iScript™ Reverse Transcription Supermix (BIO-RAD), according to the manufacturer’s instructions. For BBN Numb-WT and Numb-KO mouse BC cells, RT-qPCR was performed using the TaqMan™ Fast Advanced Cells-to-CT™ Kit (Thermo Fisher). For human bladder cancer cell lines, RT-qPCR was performed using the iQ™ SYBR® Green Supermix (BIO-RAD). The ΔCt method was used to calculate the mRNA levels of each target gene normalized against different housekeeping genes. The 2^-ΔΔCt^ method was used to compare the mRNA levels of each target gene and the relative amplification value was plotted in the graph. TaqMan Gene Expression Assay IDs were as follow: Numb (Hs01105433_m1), ANKRD1 (Hs00173317_m1; Hs00923602_g1) (Mm00496512_m1), CTGF (Hs00170014_m1; Mm01192933_g1), CYR61 (Hs00998500_g1; Mm00487498_m1), ACTB (Hs01060665_g1); GAPDH (Mm99999915_g1). The following primer sequences were used for SYBR Green methodology: GAPDH FW: 5’-GTGGAAGGGCTCATGACCA-3’, RV: 5’-GGATGCAGGGATGATGTTCT-3’; CTGF (*N.1*) FW: 5’-ACCTGTGGGATGGGCATCT-3’, RV: 5’-CAGGCGGCTCTGCTTCTCTA-3’; CTGF (*N.2*) FW: 5’-CCAATGACAACGCCTCCTG-3’, RV: 5’-TGGTGCAGCCAGAAAGCTC-3’; Cyr61 (*N1*) FW: 5’-AAATCCCCCGAACCAGTC-3’, RV: 5’-GGGCCGGTATTTCTTCACACT-3’; Cyr61 (*N2*) FW: 5’-AGCCTCGCATCCTATACAACC-3’, RV: 5’-TTCTTTCACAAGGCGGCACTC-3’; ANKRD1 (*N1*) FW: 5’-CGTGGAGGAAACCTGGATGTT-3’, RV: 5’-GTGCTGAGCAACTTATCTCGG-3’; ANKRD1 (*N2*) FW: 5’-AGCGCCCGAGATAAGTTGCT-3’, RV: 5’-CACCAGATCCATCGGCGTCT-3’.

### SDS-PAGE analysis

Cells were lysed in ice-cold RIPA buffer containing 20 mM Tris-HCl (pH 7.5), 150 mM NaCl, 1 mM Na2EDTA, 1 mM EGTA, 1% NP-40, 1% sodium deoxycholate, 2.5 mM sodium pyrophosphate, 1 mM beta-glycerophosphate, 1 mM Na_3_VO_4_, 1 µg/mL leupeptin (#9806, Cell Signaling Technology) supplemented with Halt^TM^ Protease and Phosphatase Inhibitor Cocktail (#78440, Thermo Fisher Scientific). Samples were clarified by centrifugation and protein content was measured using the BCA protein assay kit (Thermo Scientific). Proteins (10-15 μg) were resolved on 4-15% SDS-PAGE gels and transferred onto nitrocellulose membranes (Bio-Rad Laboratories). Membranes were blocked in Tris-buffered saline (TBS) containing 5% non-fat dry milk and 0.1% Tween 20 (TBS-T), prior to incubation with primary antibodies overnight at 4°C. The membranes were then washed with TBS-T followed by exposure to the appropriate horseradish peroxidase-conjugated secondary antibody for 1 hr at room temperature and visualized on X-ray film using the enhanced chemiluminescence (ECL) detection system (Bio-Rad Laboratories) or with ChemiDoc Imager followed by acquisition with Image Lab Software (Version 5.2.1).

### Rho GTPase pull-down activation assay

Activated GTP-bound RhoA was measured using the RhoA/Rac1/Cdc42 Activation Assay Combo Biochem Kit^TM^ (#BK030, Cytoskeleton, Inc, Tebu Bio Srl), following the manufacturer’s recommendations. Briefly, confluent cells were lysed in Cell Lysis Buffer (50 mM Tris pH 7.5, 10 mM MgCl2, 0.5 M NaCl and 2% Igepal) supplemented with Protease Inhibitor Cocktail (62 μg/mL Leupeptin, 62 μg/mL Pepstatin A, 14 mg/mL Benzamidine and 12 mg/mL tosyl-arginine methyl-ester) and incubated on ice for 20 minutes. Next, cell lysates were centrifuged at 13,000 rpm for 15 minutes at 4°C and supernatants were collected. For pull-down assay, 400 μg of protein lysate were added to a predetermined amount of rhotekin-RBD beads (25 μg). Twenty μg of cell lysate were used for the evaluation of total RhoA protein levels, detected using an antibody against RhoA.

### Morphometric analysis of organoids and quantitation of invasive phenotypes in 3D-Matrigel

Phase-contrast images of organoids grown in Matrigel were acquired with EVOS FL Digital Inverted Fluorescence Microscope (Invitrogen, Massachusetts) at 4x magnification after 7-12 days. Images were then imported in Fiji software^77^ (Version 2.14.0) and manually segmented to obtain regions of interest (ROI) for each organoid. Organoids smaller than 70 μm in diameter or touching the image border, and cells adherent to the well plate were excluded from the analysis. For each ROI, the analysis of organoid morphology and invasive phenotype was performed by the extraction and quantitation of the following shape descriptors: *area* (μm²), a parameter that correlates with the overall organoid size, indicative of cellular proliferation rate; *roughness* (perimeter/convex perimeter), a parameter that describes the irregularity of the organoid boundary, indicative of cell invasiveness and migratory ability; *circularity* (4π*area/perimeter²), a general measure positively correlated with organoid maturation and inversely correlated with invasiveness; *shape complexity* (number of endpoint-pixels of the skeletonized shape), a measure of the complexity of the organoid shape, indicative of aberrant growth and correlated with the number of cells protruding from the edge^32^.

### Immunofluorescence studies

For immunofluorescence (IF) staining of formalin-fixed paraffin-embedded (FFPE) samples (tissues and 3D-organoids), three µm-thick sections were deparaffinated with 100% xylene, rinsed in ethanol and re-hydrated in distilled water; antigen retrieval was performed by heating sections for 50 minutes at 95°C in pH 6.0 citrate buffer solution; tissue sections were then washed in TBS-T for 5 minutes and permeabilized with 0,3% Triton X-100 diluted in PBS for 30 minutes at room temperature (RT). Next, slides were blocked with 5% BSA diluted in PBS with 0,1% Triton X-100 for 1 hour at RT. Finally, slides were incubated overnight at 4°C with primary antibody appropriately diluted in blocking solution. The next day, sections were incubated with the proper secondary antibody and DAPI to counterstain the nuclei for 1 hour at room temperature before mounting using Mowiol solution. For the analysis of 3D-organoids, representative ROIs were acquired with a Leica SP5 Confocal Microscope. For human NMIBC TUR specimens, the whole slide was acquired with a NanoZoomer S60 Digital slide scanner (Hamamatsu Photonics K.K.) with a 20X/0.75 NA dry objective.

For the analysis of YAP on adherent cells (**Fig. 4a,e**, **6b, Extended Data Fig. 6b, 8a,b, 9a, 10b,c**), cells were co-stained for YAP and DAPI, along with any additional control markers of interest for the specific experiment and analyzed by Fiji image analysis software. The nucleus of each cell was segmented based on the DAPI channel and the mean YAP intensity was measured in an area comprising the nucleus itself and a 1 μm-thick cytoplasmic band surrounding the nucleus. The nucleus-to-cytoplasmic ratio was then computed for each cell and averaged across the different fields analyzed. For the analysis of YAP in mouse FFPE samples (**Fig. 3a**, **Extended Data Fig. 4b-d**), sections were co-stained for active (non-phosphorylated) YAP and DAPI. The nucleus of each cell was segmented based on the DAPI channel in Fiji followed by quantification of the mean active-YAP nuclear intensity. Cells with a nuclear intensity higher than 50 (arbitrary unit) were classified as YAP-positive.

For the analysis of YAP in FFPE human NMIBC TUR samples (Fig. 1j), FFPE sections were co-stained for active YAP and DAPI and analyzed by QuPath (v. 0.4.4)^78^. Nuclei were automatically segmented on the DAPI channel using the deep-learning Stardist algorithm, based on an in-house trained model. The cytoplasmic cell area was represented as a 3 µm-thick band around the nucleus. Intensity of active YAP staining was measured in both nuclear and cytoplasmic cell compartments. For each patient, three independent representative regions of tumor tissue, each of average area equal to 1 mm^2^, were manually selected and analyzed for the distribution of cells with negative (nuclear intensity comparable to background cytoplasmic intensity) or positive (nuclear intensity 1.5-fold superior to background cytoplasmic intensity) active YAP nuclear expression. Detection measurements relative to each region were exported in R Studio (v. 2023.06.2). Filters were applied on the exported database to exclude non-reliable cell detections (objects with size lower than 15 µm^2^ and intensity lower than a custom threshold in the DAPI channel were discarded). The number of active YAP-positive cells was then normalized to the total number of cells per each region and aggregated per patient to extract the mean percentage of positive cells.

The quantitative analysis of pMLC2 and pCofilin staining (Fig. 5b, Extended Data Fig. 9b,c) was conducted in Fiji by computing the mean fluorescence intensity over the entire area of the field of view occupied by cells. The segmentation of this area was achieved by generating an image from the maximum intensity projection of all channels, following automatic brightness and contrast adjustment to scale the signals. A custom threshold was then established on this image to delineate the cell-occupied regions for intensity measurement.

### Immunohistochemistry

Three-μm thick sections were prepared from FFPE tissue blocks, dried at 37°C O/N and then processed with Bond-RX fully automated stainer system (Leica Biosystems) according to the following protocol. For phospho-Cofilin and Numb staining, antigen retrieval was performed by heating sections in a microwave oven for 10 minutes on high power (∼900 watt) in citrate buffer solution pH 6.0 at 100°C for 40 min. After the washing steps, peroxidase blocking was performed for 10 min using the Bond Polymer Refine Detection Kit (#DC9800, Leica Biosystems). Tissues were incubated for 30 min with the appropriate Ab diluted in Bond Primary Ab Diluent (#AR9352). Subsequently, tissues were incubated with post primary and polymer for 16 min, developed with DAB-chromogen for 10 min and counterstained with hematoxylin for 5 min. Slides were digitally scanned with the Aperio ScanScope. Digital images were processed with Adobe Photoshop CS3 (Adobe Inc.).

### Transwell Invasion assay

Transwell migration/invasion assays were performed using PET membrane inserts (BRAND® 24-well Cell Culture Insert, PET-membrane, #BR782711, Merck) covered with 20 μL of Matrigel:PBS (1:2), as described in the manufacturer’s protocol. Briefly, 1×10^5^ cells were seeded in growth factor-or serum-free medium, depending on the cell lines, in the upper chamber of the transwell insert, and treated, when necessary, with the appropriate pharmacological inhibitors (added in the upper and lower chamber of the transwell). Complete medium supplemented with HGF (25 ng/mL) was added to the lower part of the transwell. After an appropriate incubation period (18 hr for BBN-induced mouse tumor cells, 24 hr for RT112 cells, 48 hr for RT4 cells), the transwell inserts were removed and the cells were fixed in the transwell insert with 4% PFA for 15 minutes. Cells were then washed with water to remove the formaldehyde. Then, using a sterile cotton swap, cells which had not migrated through the membrane were scraped off the top of the transwell insert and stained by using DAPI+0.1% Triton X-100 in PBS for 30 minutes. Images corresponding to the entire area of the transwell were collected with the Nikon Ti-2 microscope using a 10x/0.3 NA dry objective and the number of invaded cells was analyzed using the Fiji software.

### RNA sequencing

Libraries were prepared using the TruSeq Stranded Total RNA Gold kit (Illumina) according to the manufacturer’s protocol. Sequencing was performed on an Illumina NovaSeq 6000 platform with paired-end reads of 50 bp. Sequenced raw reads were processed with NF-CORE RNASeq pipeline (https://github.com/nf-core/rnaseq, version 3.8.1)^79^, using the human GRCh38 and mouse GRCm39 as reference genomes. Transcript abundances were quantified using the Salmon pseudo-aligner^80^. Genes abundance normalization and differential expression testing were performed with the DESeq2 R package (version 1.34)^81^. Log2 fold changes obtained were shrunken using the "apeglm" method^82^ to improve the stability of the estimates in the presence of a limited number of replicates. In the absence of replicates for the BBN-KO sample treated with verteporfin *vs.* vehicle, Log2 fold changes were calculated by directly subtracting log1p(normalized counts) of each condition, considering only genes with an average normalized count >20. All samples for each condition tested were analyzed with principal component analysis and projected. When the batch effect was associated with a principal component that explained more than 20% of the variance, the batch of the experiment was added as a covariate in the statistical model. Genes with a Benjamini-Hochberg adjusted p-value <0.05 and an absolute log2 fold change >0.5 were uploaded to the Ingenuity Pathway Analysis (IPA) software (Qiagen). IPA was used to infer upstream regulators from the RNAseq data by applying the "Upstream Regulator Analysis" tool^83^. Transcription regulators with an activation Z-score >1.5 or <-1.5 and an overlap p-value <0.05 were considered as significantly activated or inhibited, respectively. The association between gene expression modifications and changes in pathways and biological states was assessed with a Gene Set Enrichment Analysis (GSEA)^6^. The analysis was performed with the “fgsea” method implemented in the ClusterProfiler R package (version 4.2.2)^84^ and the MSigDB “Hallmarks”^85^ set was used as annotation. Genes were ranked based on their log2 fold change values. Genes with low counts or high dispersion that did not pass the DESeq2 independent filtering were not considered for the analysis. For the evaluation of the activity of YAP/TAZ pathway (not included in the “Hallmarks” set), a signature composed of 22 direct YAP/YAZ transcriptional targets^16^ was additionally used as annotation for the GSEA.

### Definition of the NUMB^LESS^ signature and clinical validation of its prognostic relevance to NMIBC

We established a gene signature reflective of Numb deficiency in human bladder cancer by identifying 15 consistently upregulated genes (Log2FC >1, padj <0.01) and 12 consistently downregulated genes (Log2FC <-1, padj <0.01) following Numb knockdown in all three Numb-proficient bladder cancer cell lines (RTT4, RT112, KK47). Raw gene abundances from bladder cancer cell lines with high and low endogenous Numb expression, alongside abundances from Numb-KD experiments, were normalized using DESeq2. Normalized abundances were then converted to a log scale with the variance stabilizing transformations (implemented in DESeq2), before z-score computation. Using the expression levels of these 27 signature genes, we conducted unsupervised clustering of the samples with the Hierarchical Clustering algorithm from the complexHeatmap R library (version 2.13.4), employing a 1-Pearson’s correlation as the distance metric. We then utilized this 27-gene signature to cluster patients from the 535 NMIBC patients of the UROMOL cohort^20^. Raw gene abundances were normalized, transformed, and scaled as mentioned above. We then applied the Partitioning Around Medoids (PAM) algorithm for clustering, using the 1-Pearson’s correlation coefficient as the distance metric. The silhouette method used to determine the optimal number of clusters identified two distinct NMIBC tumor subtypes. The 27 genes in the signature were hierarchically clustered to confirm the consistent and coordinated modulation of genes upregulated and downregulated in clusters predicted as NUMB^LESS^ “like” and “not-like”. The 22-gene YAP/TAZ transcriptional target signature was used to classify the NMIBC samples into YAP-active and YAP-inactive groups. The PAM algorithm was reapplied for clustering with the 1-Pearson’s distance metric. The differential propensity for MIBC progression among the two NMIBC subtypes, as defined by the NUMB^LESS^ signature, was assessed using a univariable and multivariable Cox proportional hazards model. Multivariable analyses were adjusted for clinically relevant variables (age, gender, tumor grade, pT, history of intravesical therapy).

### Statistical analysis and reproducibility

PCR experiments were performed in biological triplicates with the exception of the control PCR experiments of **Extended Data Fig. 9d** in a single replicate. All the other experiments were repeated a minimum of two times. The number (n) of cells, animals, fields, patients, and independent experiments is reported in figures or in accompanying figure legends. Data are represented as mean ± standard error (SEM) in all cases except for Fig. 3a, and Circularity, Roughness, and Shape Complexity measures in **Fig. 3c-d**, **4g**, **5f**, **6f**, **Extended Data Fig. 5c, 6d, 8d, 10e** which were represented as boxplots. The boxplot is depicted with a box covering the interquartile range (IQR) from the 25th to the 75th percentile, the whiskers extend from the ends of the box to the smallest and largest data values within a 1.5 IQR. The horizontal line indicates the median, and the “X” indicates the mean. Means were compared using two sample Welch’s t-test, except for comparisons with the reference sample in RT-qPCR experiments (**Fig. 3g**, **4c**, **5d**, **6e**, **Extended Data Fig. 6c**, **10d**), where one-sample t-test was performed. When multiple conditions were present we performed a FDR adjusted pairwise Welch’s t-test following a significant Welch’s ANOVA except for **Fig. 4a**,**f**, **5c**, **6b**, **Extended Data Fig. 4b,c, 8b**, **10c** which were analyzed with Tukey HSD test following a significant ANOVA. Data distribution was assumed to be normal although not formally tested. The Benjamini-Hochberg method was employed for FDR corrections in multiple comparison testing. When performing multiple comparisons, only comparisons of biological interest are shown in the graphs; complete results are available in the source data.

Differences in proportions were tested with Fisher’s exact t-test in all 2x2 tables (**Fig. 1i**, **Extended Data Fig. 1a,c**) and with Pearson’s Chi-squared test in in larger contingency tables. (**Fig. 2b,c**). Odds ratio and relative confidence intervals were calculated with Small Sample-Adjusted Unrestricted Maximum Likelihood Estimation and Normal Approximation (Wald) for **Fig. 2b,c** and with the Median-Unbiased Estimate and Mid-P Exact Confidence Intervals for **Extended Data Fig. 1a,c.** For survival analysis, the Cox proportional hazards model was employed as implemented in the ’coxph’ function of the survival R package (version 3.5) to compute hazard ratios (HR) and confidence intervals (CI) in both univariable and multivariable analyses. Global significance was tested using the log-rank test, while the significance of each variable was determined using the Wald test. Statistical tests were two-sided with significance set at p < 0.05. Significance in figures is indicated as: ∗, p < 0.05; **, p < 0.01; ***, p < 0.001; ****, p < 0.0001; ns, not significant. Exact p-values are reported in source data. All statistical analyses were performed using the R software (version 4.1.2). Details of each test are provided in the figure legends.

## Data availability

The raw and processed RNA sequencing data generated in this study have been deposited in the GEO database. The clinicopathological data from the UROMOL cohort used in this study are available in the source data of the referenced publication. Raw gene abundances were kindly provided by the authors. Human and mouse gene set annotations for pathway analysis were retrieved from MSigDB [https://www.gsea-msigdb.org/gsea/msigdb/] with the msigdbr package (version 7.4.1) in R. Human GRCh38 and mouse GRCm39 reference genomes (primary genome assembly) used for the alignment of RNA sequencing data are available on GENCODE [https://www.gencodegenes.org/]. All data are available in the main text or the source data provided with this paper.

## ACKNOWLEDGMENTS

We thank the anonymous patients who donated their samples for research and the IEO Pharmacy for chemotherapy drugs. We thank R. Gunby for critically reading the manuscript; A. Gobbi, M. Capillo, and the Mouse facility, and the Real Time PCR and DNA Sequencing Service of Cogentech (Cogentech Srl, Milan) for mice handling and technical support; L. Rotta and the IEO Genomic Unit for RNA sequencing experiments; G. Bertalot, Luise C., Ricca D., Tillhon M., F. Montani and the IEO Molecular and Digital Pathology Unit for technical assistance with tissue processing, staining and image acquisition; D. Disalvatore for help with statistical analyses; M. Monturano, G. Peruzzotti, the IEO Clinical Trial Office and the Retrospective Data Governance Board for assistance with clinical databases and ethical procedures; O. De Cobelli, F. Nolé, A. Mistretta, V.D. Matei, G. Cordima, G. Cozzi and the IEO Urology Program for help with clinical cohorts. M.G.F. was supported by a fellowship from the Fondazione Umberto Veronesi (FUV), R.P. and D.C.R. were supported by an AIRC fellowship. This work was supported by: Associazione Italiana per la Ricerca sul Cancro (AIRC-IG 23049 to S.P., IG 23060 to P.P.D.F.); PSR-2020/2022 from the University of Milan to S.P.; The Italian Ministry of University and Scientific Research (MIUR-PRIN2017 and MIUR/PRIN2020 to S.P); The Italian Ministry of Health, RF-2016-02361540 e RF-2021-12373957 to P.P.D.F.; The Italian Ministry of Health with Ricerca Corrente and 5x1000 funds.

## AUTHOR CONTRIBUTIONS

Conceptualization: D.T. and S.P.

Methodology: R.P., D.C.R., M.G.F., R.B., F.R. performed the in vitro experiments; D.T. and R.B. performed the ex vivo experiments; F.A.T., F.S., G.R., N.F. and G.V. performed pathological analyses; G.J. and R.B. did IHC stainings.

Formal analysis: F.A.T., C.S. and S.R., imaging analysis; F.A.T., bioinformatic and statistical analysis on gene expression; F.A.T. and S.F. statistical analysis of patient data.

Resources: F.S., G.R., N.F., P.P.D.F., G.V., G.P., G.M., S.P.

Data curation: F.A.T., R.P, D.C.R., M.G.F., S.F, F.R.

Investigation: F.A.T., R.P., D.C.R., M.G.F., D.T.

Writing the original draft: D.T., S.P., F.A.T.

Writing (review and editing): All Authors.

Supervision: S.P. and D.T.

Funding acquisition: S.P., D.T., P.P.D.F., G.P.

D.T. and S.P. are the custodians of all the original documentation.

## DATA AND MATERIALS AVAILABILITY

All data are available in the main text or the supplementary materials. Correspondence and material requests should be addressed to.

