## Extended Data for "Loss of the tumor suppressor NUMB drives aggressive bladder cancer through hyperactivation of a RhoA/ROCK/YAP signaling circuitry"

**EXTENDED DATA FIGURES AND LEGENDS**

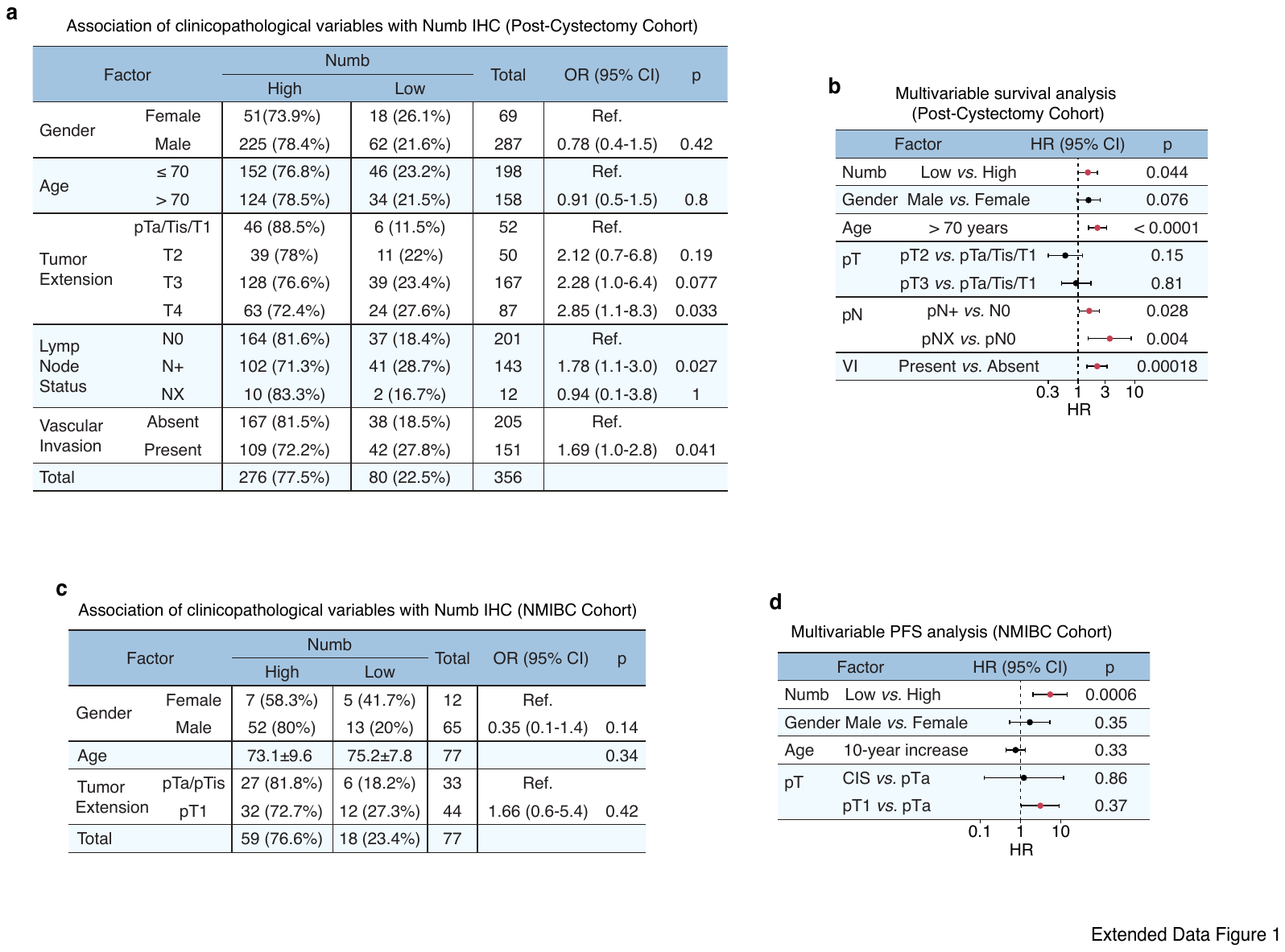

### Extended Data Figure 1. Numb is a prognostic biomarker of unfavorable disease course in real-life MIBC and NMIBC patients. a. Association between clinicopathological variables and Numb IHC status (High *vs.* Low) in the retrospective longitudinal cohort of 356 post-cystectomy MIBC patients. OR (95% CI), Odds Ratio (95% Confidence Interval), in this and other relevant panels in the figure; p, Fisher’s exact test p-value. b. Multivariable survival analysis of the association between the indicated factors and good (HR<1) or poor (HR>1) prognosis in 258 patients with available follow-up from the MIBC cohort in ‘a’, by Cox proportional hazards model. Significant associations are marked in red. HR, multivariable hazard ratios, in this and other relevant panels in the figure; p, Wald-test p-value. c. Association between clinicopathological variables and Numb IHC status (High *vs.* Low) in a retrospective longitudinal cohort of 77 NMIBC patients; p, Fisher’s exact test p-value for categorical variables and Welch’s t-test for Age. d. Multivariable progression-free survival (PFS) analysis of the association between the indicated factors and good (HR<1) or poor (HR>1) prognosis in the 77 NMIBC patient cohort from ‘c’, by Cox proportional hazards model; p, Wald-test p-value.

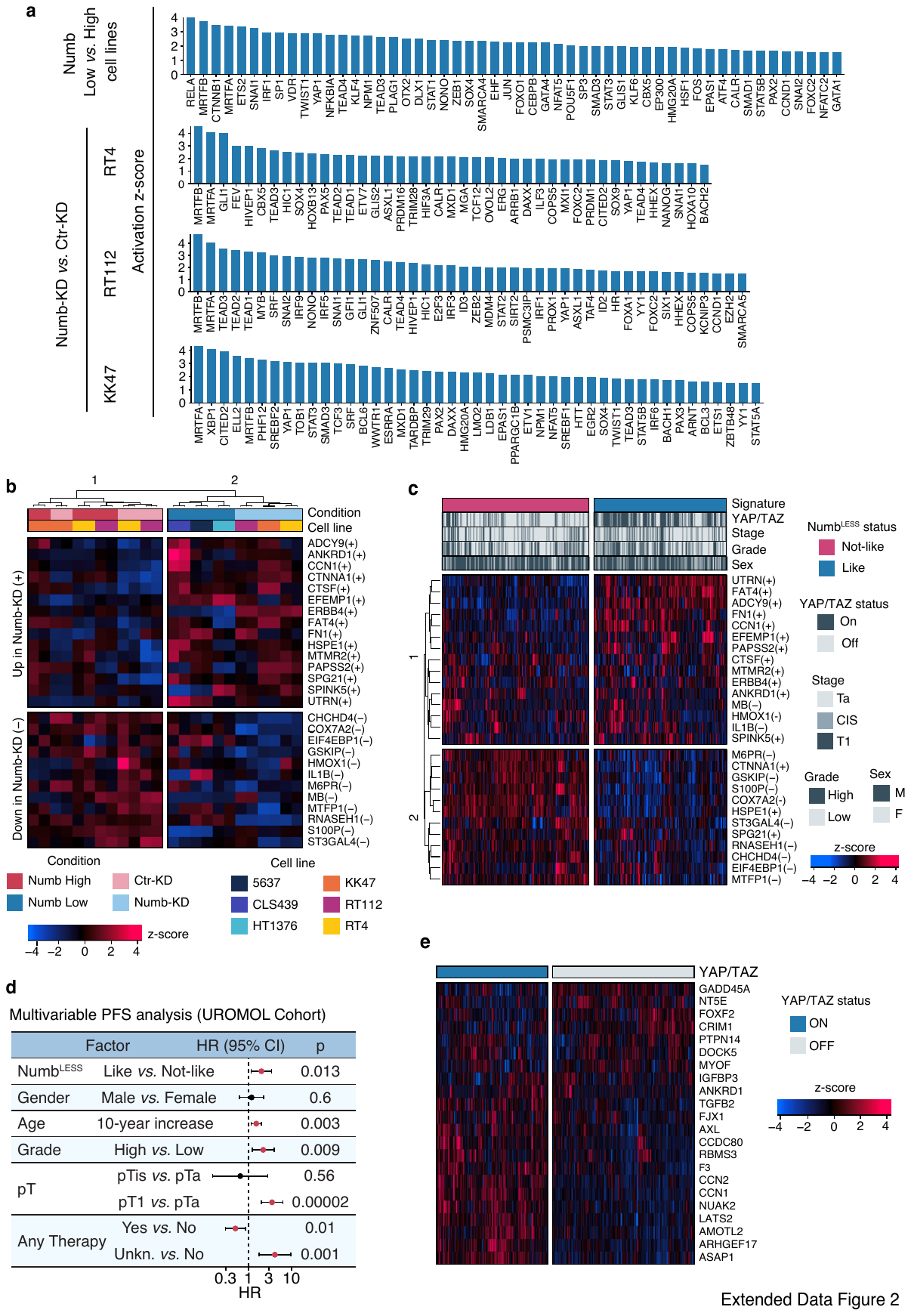

### Extended Data Figure 2. Validation of a multigene signature informative of a Numb-deficient condition in preclinical human BCa cell models and in real-life NMIBC patients. a. Transcriptional regulators predicted to be activated in MIBC Numb^Low^ (HT1376, CLS439, 5637) *vs.* NMIBC Numb^High^ BCa (KK47, RT4, RT112) cell lines (top) and in RT4, RT112 and KK47 Numb-KD *vs.* Ctr-KD cells (bottom). p-value of overlap <0.05 by Fisher’s exact test. Activation z-score>1.5. b. Hierarchical unsupervised clustering of Numb^High^ and Numb^Low^, and Numb-KD and Ctr-KD BCa cell lines as described in ‘a’, based on the expression levels of 27 genes consistently upregulated (+) or downregulated (-) (FDR adjusted p-value<0.01, Log2FC >1 or <-1) in Numb-KD *vs.* Ctr-KD KK47, RT4, RT112 cells (Numb^LESS^ signature). The color code scale indicates the z-score of log-normalized transcript abundance (ranging from -4 to 4), in this and other relevant panels in the figure. Genes are shown on the right, cell lines and conditions are indicated on the top and bottom, and direction of gene regulation on the left. Groups are indicated on the top. c. Heatmap showing the unsupervised clustering of 535 NMIBC patients from the UROMOL cohort, stratified by the Numb^LESS^ signature (Numb^LESS^-Like *vs*. -Not-Like). Upregulated (+) or downregulated (-) genes comprised in the Numb^LESS^ signature are indicated on the right, together with color codes for YAP/TAZ activation status, stage, grade and sex. Groups are indicated on the top (Numb^LESS^-Like and -Not-Like) and left (groups 1 and 2, indicating clusters of genes with consistent and coordinated up- or down-modulation in Numb^LESS^-Like and -Not-Like groups). d. Multivariable progression-free survival (PFS) analysis of the association between the indicated factors and good (HR<1) or poor (HR>1) prognosis in 535 NMIBC patients from the UROMOL cohort, by Cox proportional hazards model. Significant associations are marked in red. HR, multivariable hazard ratios; 95% confidence intervals (CI); p, Wald-test p-value. e. Heatmap showing the unsupervised clustering of 535 NMIBC tumors from the UROMOL cohort, classified as YAP/TAZ active (ON) or inactive (OFF), based on the expression levels of 22 genes of the YAP/TAZ signature.

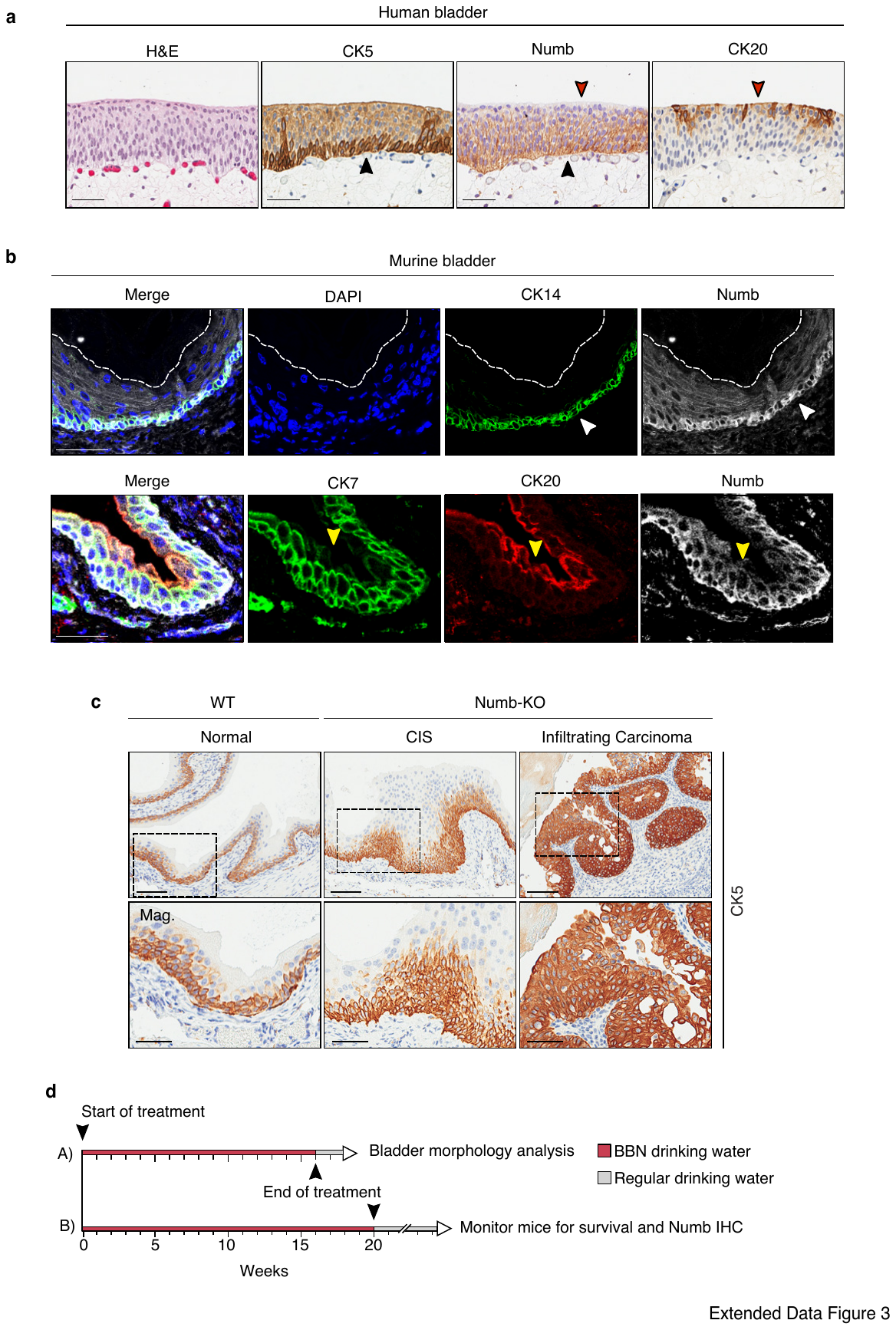

### Extended Data Figure 3. Characterization of the pro-tumorigenic role of Numb loss in the Numb-KO mouse model. a. Representative IHC images of the expression of endogenous Numb in the normal human urothelium, showing its association with the basal layer. CK5 is a marker for the basal/myoepithelial layer (black arrowheads); CK20 is a marker for superficial cells (red arrowheads). Bars, 50 µm. b. Representative confocal fluorescence images of FFPE sections of murine bladder. Upper panels show co-staining for endogenous Numb (white), the CK14 basal/myoepithelial layer marker (green) and DAPI (blue). White arrowheads indicate the basal layer. Lower panels show co-staining for Numb (white), the intermediate layer marker CK7 (green), and the superficial/umbrella cell marker CK20 (yellow arrowheads). The dashed line delineates the superficial layer of the usothelium. Bars, 50 µm. c. Upper panels, representative IHC images of CK5 expression in bladder tissue from Numb-KO and WT mice showing expansion of basal cells (CK5+) in Numb-KO lesions (CIS and infiltrating cancer) compared with WT. Boxed areas are magnified (Mag.) in the lower panels. Bars, 100 μm; Mag., 50 μm. d. BBN treatment protocols for WT *vs.* Numb-KO mice. A. For the analysis of the incidence of hyperplastic, not-infiltrating CIS and infiltrating cancer lesions in the WT *vs.* Numb-KO bladder mucosa (shown in Fig. 2c), mice were administered 0.05% BBN in drinking water continuously for 16 weeks and then switched to regular drinking water. Mice were sacrificed two weeks after ending BBN treatment. B. For survival and histological analyses (shown in Fig. 2d-f), mice were administered 0.05% BBN in drinking water continuously for 20 weeks to induce the formation of infiltrating tumors, and then switched to regular drinking water. Mice were monitored for survival until the experimental endpoint, at which time all surviving mice were sacrificed. Bladder tissues were subjected to IHC analysis of Numb expression.

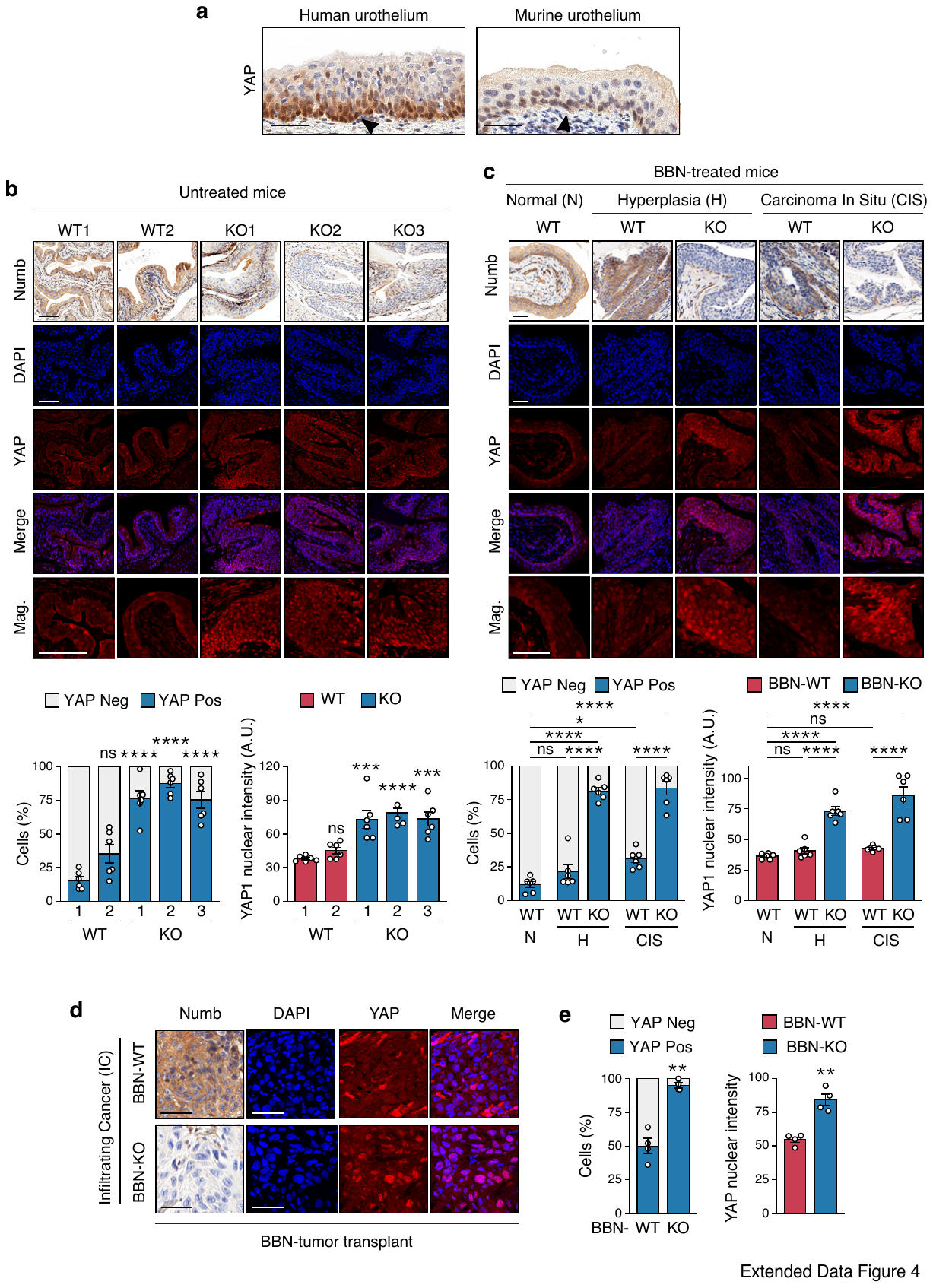

### Extended Data Figure 4. Numb loss is associated with increased nuclear accumulation of YAP in the normal, preneoplastic and overtly neoplastic urothelium. a. Representative IHC images showing the expression pattern of YAP in the normal human and murine urothelium. The black arrowheads point to the basal layer. Bars, 50 µm. b. Top, Representative images of bladder tissues obtained from treatment-naïve adult WT (n=2; WT1, WT2) and Numb-KO (n=3; KO1, KO2, KO3) mice, examined by IHC for Numb expression and co-stained for YAP (red) and DAPI (blue) by immunofluorescence (IF). Merged images of YAP and DAPI staining and magnifications (Mag.) are shown. Bars, 100 μm. Bottom, Quantification of the % of YAP-positive cells and mean nuclear YAP intensity expressed as mean ± SEM, n=6 fields for each condition. ****, p<0.0001; ***, p<0.001; ns, not significant, *vs.* WT1, by Tukey's HSD test. c. Top, Bladder tissues from 16-week old BBN-treated WT and Numb-KO mice were examined by IHC for Numb and co-stained for YAP (red) and DAPI (blue) by IF. Shown are representative images of normal urothelium (N), hyperplasia (H) and carcinoma *in situ* (CIS) detected in a WT bladder, and of hyperplasia (H) and CIS detected in a Numb-KO bladder. Bars, 50 μm. Bottom, Quantification of the % of YAP-positive cells and mean nuclear YAP intensity expressed as mean ± SEM, n=6 fields for each condition. ****, p<0.0001; *, p<0.05; ns, not significant, by Tukey's HSD test. d. Primary infiltrating cancers induced by 20 weeks of BBN treatment in WT and Numb-KO mice as in ‘c’ were excised and transplanted into NSG mice. Then, secondary WT and Numb-KO BBN-tumor transplants (BBN-WT and BBN-KO) were excised and examined by IHC for Numb and co-stained for YAP (red) and DAPI (blue) by IF. Bars, 50 μm. e. Quantification of the experiment in ‘d’. Graphs show the % of YAP-positive cells and mean nuclear YAP intensity expressed as mean ± SEM, n=4 fields for each condition. **, p < 0.01; by Welch’s t-test *vs.* WT.

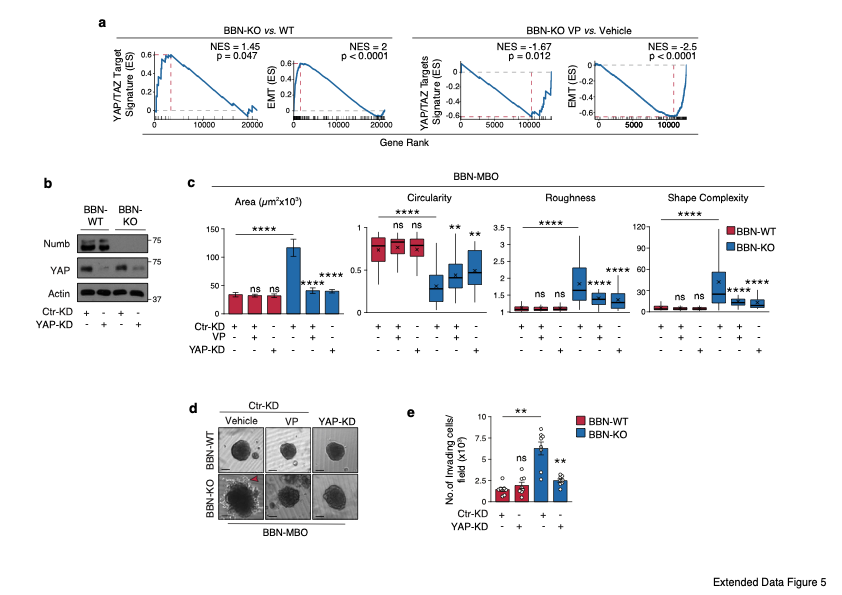

### Extended Data Figure 5. Aberrant morphogenesis and invasiveness of BBN-KO tumor cells are dependent on YAP. a. GSEA enrichment plot of the YAP/TAZ and EMT gene signatures in RNA-seq data from BBN-KO *vs.* BBN-WT tumor cells (left, n=3 for each condition) and BBN-KO cells treated with VP *vs.* vehicle (right, n=1 for each condition). ES, Enrichment Score; NES, Normalized Enrichment Score; p, permutation test p-value. b. BBN-WT and BBN-KO cells were lentivirally transduced with a shRNA targeting luciferase as a control (Ctr-KD) or YAP (YAP-KD). Immunoblot analysis of YAP silencing efficiency is shown. Actin, loading control. Results are representative of two independent experiments. c. BBN-WT and BBN-KO cells transduced as in ‘b’ were cultured as 3D-organoids (BBN-WT and BBN-KO MBO) in Matrigel. Ctr-KD BBN-WT and BBN-KO cells were also treated with vehicle or 25 nM Verteporfin (VP) as indicated. Graphs show the quantitation of the indicated morphometric parameters. Area (μm2) is reported as mean ± SEM. Other parameters are reported as boxplots delimited by 25th and 75th percentiles and showing the median (horizontal line) and the mean (X). The whiskers span from the smallest and largest data values within a 1.5 interquartile range. Data were obtained from two independent experiments. ****, p<0.0001; **, p<0.01; ns, not significant *vs.* matching condition, by FDR-adjusted pairwise Welch's t-test. d. Representative bright field images of BBN-MBO from the experiment in ‘c’. Bar, 100 μm. The red arrowhead indicates invading protrusions. e. Transwell Matrigel invasion assay of BBN-WT and BBN-KO cells, control-silenced (Ctr-KD) or YAP-silenced (YAP-KD). Graph shows the number of invading cells/field 24 h after seeding, expressed as the mean ± SEM of 8 fields from two independent experiments. **, p<0.01; ns, not significant, *vs*. matching controls by FDR-adjusted pairwise Welch's t-test.

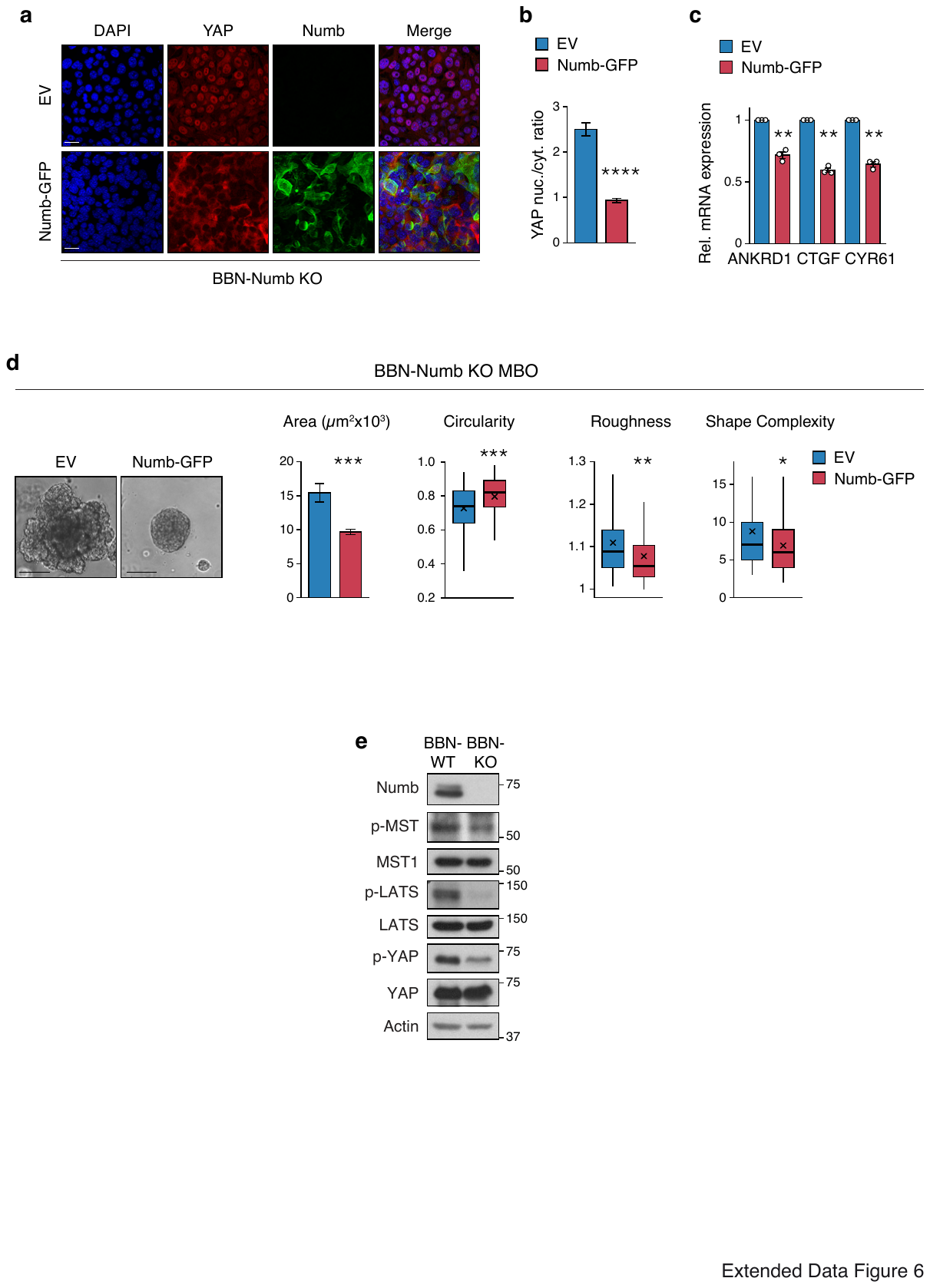

### Extended Data Figure 6. Restoration of Numb expression in BBN-KO tumors reverses YAP nuclear accumulation and invasive phenotypes. a. Representative confocal images of BBN-KO tumor cells stably infected with a Numb-GFP lentiviral vector (Numb-GFP) or an empty vector (EV) and co-stained for endogenous YAP by IF (red) and DAPI (blue). Bar, 25 μm. b. Quantification of YAP nuclear/cytoplasmic ratio in cells described in ‘a’. Results are reported as the mean ratio/field ± SEM, n=12 fields for each condition from three independent experiments. ****, p<0.0001 by Welch’s t-test *vs.* EV. c. RT-qPCR analysis of the indicated YAP transcriptional targets in BBN-KO tumor cells transduced with Numb-GFP as in ‘a’. Graphs show the relative mean fold expression ± SEM from three independent experiments. **, p<0.01 by one-sample t-test. d. Representative bright field images and quantitation of morphometric parameters of 3D-Matrigel MBO generated by BBN-KO tumor cells (BBN-Numb KO MBO), transduced as in ‘a’. EV, n=79; Numb-GFP, n=123, from three independent experiments. Area (μm^2^) is reported as mean ± SEM. Other parameters are reported as boxplots delimited by 25^th^ and 75^th^ percentiles and showing the median (horizontal line) and the mean (X). The whiskers span from the smallest and largest data values within a 1.5 interquartile range. ***, p<0.001; **, p<0.01; *, p<0.05, relative to matching condition, by FDR-adjusted pairwise Welch's t-test *vs.* EV. e. Immunoblot analysis of the total and phosphorylated levels of the indicated Hippo pathway components (MST1/2, LATS, YAP) in BBN-WT *vs.* BBN-KO cells. Numb levels are shown. Actin, loading control. Blot is representative of three independent experiments.

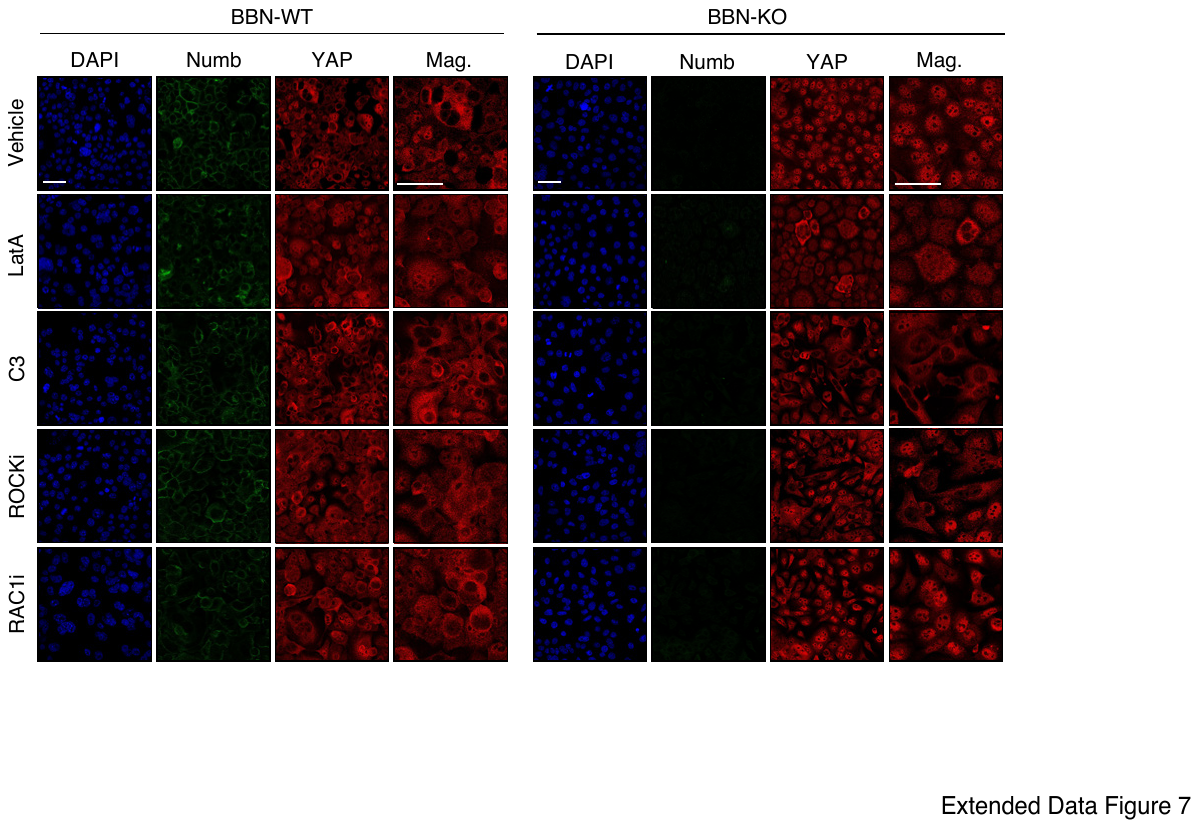

### Extended Data Figure 7. Effects of RhoA, Rac1, ROCK and actin remodeling inhibition on YAP nuclear translocation in BBN-WT and BBN-KO tumor cells. Representative confocal fluorescence images of BBN-WT and BBN-KO tumor cells treated with the indicated inhibitors (see also Legend to Fig. 4a) and co-stained for endogenous YAP (red), Numb (green) and DAPI (blue). A magnification (Mag.) for each condition is shown. Bars, 50 µm.

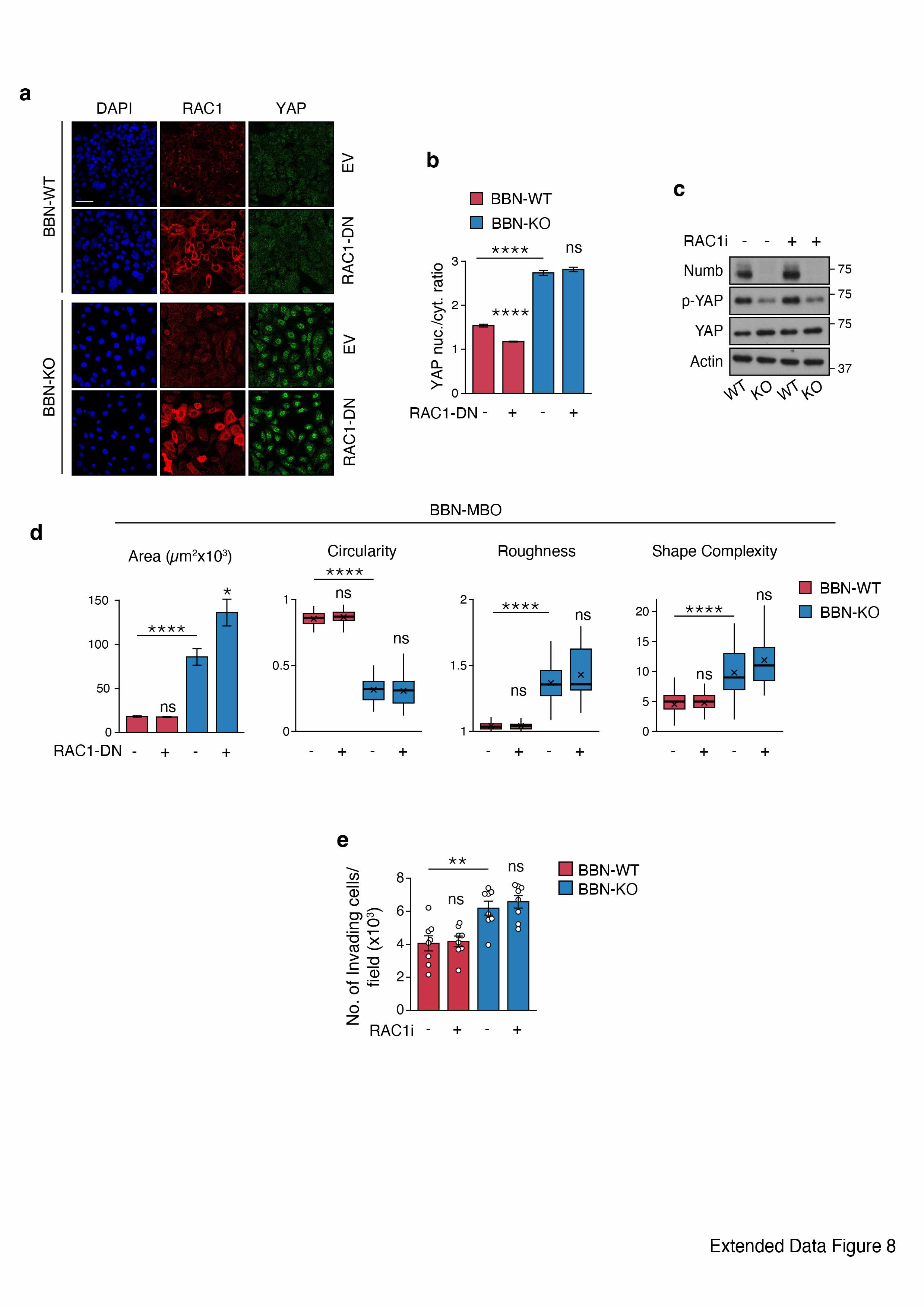

### Extended Data Figure 8. Rac1 inhibition does not affect YAP hyperactivation induced by loss of Numb. a. Representative confocal images of BBN-WT and BBN-KO tumor cells transduced with a lentiviral vector encoding a dominant negative Rac1 mutant (RAC1-DN) or empty vector (EV) and co-stained for endogenous Rac1 (red), YAP (green) and DAPI (blue). Bar, 50 µm. b. Quantification of YAP nuclear/cytoplasmic ratio in cells described in ‘a’. Graphs show the mean ± SEM, n=14 fields, from two independent experiments. ****, p<0.0001; ns, not significant, *vs.* matching controls by Tukey's HSD test. c. Immunoblot analysis of Numb and total and phosphorylated YAP in BBN-WT and BBN-KO cells treated with the Rac1 inhibitor NSC-23766 (RACi, 10 µM for 12 h). Actin, loading control. Blots shown are representative of two independent experiments. d. Analysis of morphometric parameters of BBN-MBO derived from cells treated as described in ‘a’. Area (μm^2^) is reported as mean ± SEM. Other parameters are reported as boxplots delimited by 25^th^ and 75^th^ percentiles and showing the median (horizontal line) and the mean (X). The whiskers span from the smallest and largest data values within a 1.5 interquartile range. Data were obtained from two independent experiments. ****, p<0.0001; ns, not significant, relative to matching condition by FDR-adjusted pairwise Welch's t-test. e. Transwell Matrigel invasion assay of BBN-WT and BBN-KO cells treated with the Rac1 inhibitor, NSC-23766 (RACi, 250 nM for 24 h). Graph shows the average number of invading cells/field calculated 24 h after seeding, expressed as the mean ± SEM of 8 fields from two independent experiments. **, p<0.01; ns, not significant, *vs*. matching controls, by FDR-adjusted pairwise Welch's t-test.

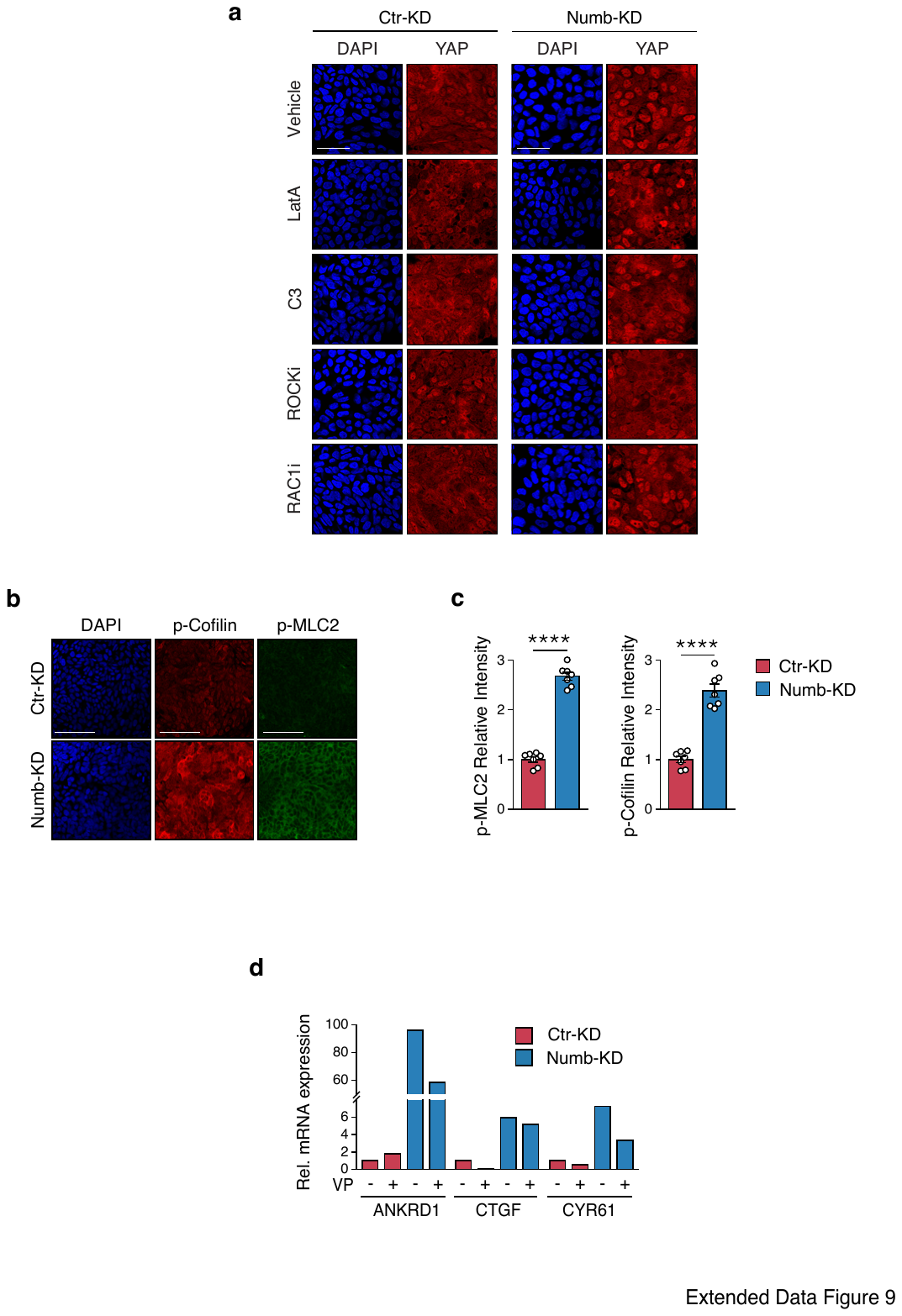

### Extended Data Figure 9. Numb silencing induces increased RhoA/ROCK activity and YAP hyperactivation in human RT4 BCa cells. a. Representative confocal images of Ctr-KD and Numb-KD RT4 cells treated with the indicated inhibitors (see Legend to Fig. 6b) and co-stained for endogenous YAP (red) and DAPI (blue). Bar, 50 µm. b. Representative confocal images of Ctr-KD and Numb-KD RT4 cells co-stained for endogenous phosphorylated cofilin (p-Cofilin, red), phosphorylated MLC2 (p-MLC2, green) and DAPI (blue). Bar, 50 µm. c. Quantification of the experiment in ‘b’ showing the relative intensity of p-MLC2 (left) and p-Cofilin (right) in each condition. Values are the mean/field ± SEM, n=6 fields/condition from one experiment, representative of two independent experiments. ****, p<0.0001 by Welch’s t-test *vs.* Ctr-KD. d. Representative RT-qPCR analysis for the indicated YAP transcriptional targets in Numb-KD and Ctr-KD RT4 cells treated with Verteporfin (VP, 3 µM for 8 h) or vehicle-treated. Results are reported as the relative mean fold expression from one experiment run in triplicate.

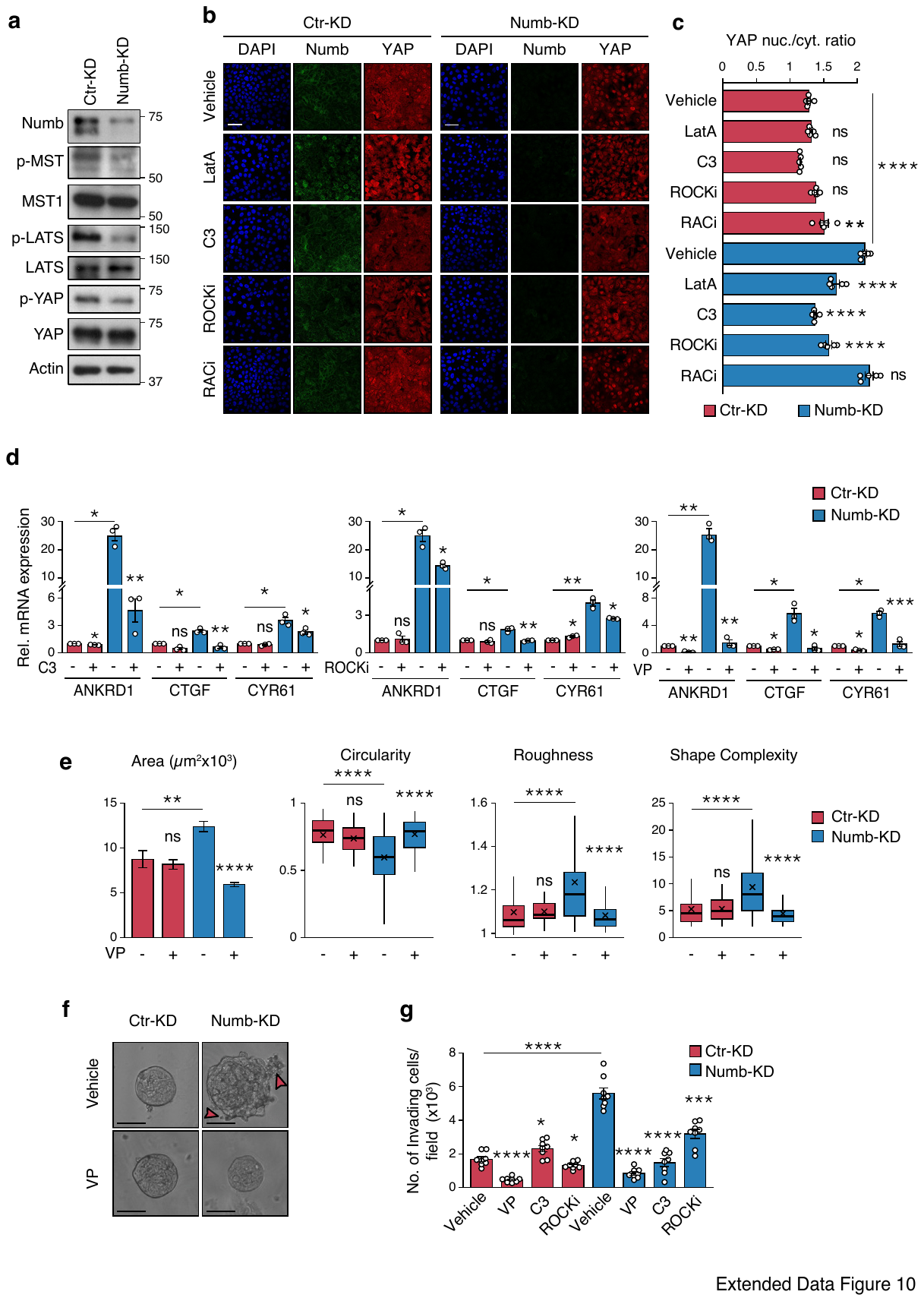

### Extended Data Figure 10. Numb silencing induces RhoA/ROCK-dependent YAP hyperactivation in human RT112 BCa cells. a. Immunoblot analysis of Numb and of the total and phosphorylated levels of the indicated Hippo pathway components (MST1/2, LATS and YAP) in Ctr-KD and Numb-KD RT112 cells. Actin, loading control. Blots are representative of two independent experiments. b. Representative confocal images of Ctr-KD and Numb-KD RT112 cells treated with the actin polymerization inhibitor Latrunculin-A (LatA, 500 nM for 6 h), RhoA inhibitor C3 transferase (C3, 3 µg/ml for 6 h), ROCK inhibitor Y-27632 (ROCKi, 10 µM for 12 h), Rac1 inhibitor, NSC-23766 (RACi, 10 µM for 12 h), or vehicle, and co-stained for endogenous YAP (red), Numb (green) and DAPI (blue). Bar, 50 µm. c. Quantification of the experiment in ‘b’ showing YAP nuclear/cytoplasmic ratio in the different conditions. Values are the mean/field ± SEM, n=5 fields/condition from one experiment, representative of two independent experiments. ****, p<0.0001; **, p<0.01; ns, not significant, relative to matching controls by Tukey’s HSD test. d. Representative RT-qPCR analysis for the indicated YAP transcriptional targets in Ctr-KD *vs.* Numb-KD RT112 cells treated with C3 (3 µg/ml for 6 h) (left), ROCKi (50 µM for 8 h) (middle), Verteporfin (VP, 3 µM for 8 h) (right), or vehicle-treated. Graphs show the relative mean fold expression ± SEM from three independent experiments. ***, p<0.001; **, p<0.01; *, p<0.05; ns, not significant, *vs.* matching condition, by FDR-adjusted unpaired one-sample t-test (*vs.* reference sample) or two-sample Welch’s t-test (treated *vs.* vehicle Numb-KD samples). e. Analysis of morphometric parameters in 3D-Matrigel organoids derived from Ctr-KD and Numb-KD RT112 cells treated with VP (25 nM) or vehicle. Area (μm^2^) is reported as mean ± SEM. Other parameters are reported as boxplots delimited by 25^th^ and 75^th^ percentiles and showing the median (horizontal line) and the mean (X). The whiskers span from the smallest and largest data values within a 1.5 interquartile range. Data were obtained from two independent experiments. ****, p<0.0001; **, p<0.001; ns, not significant, relative to matching condition by FDR-adjusted pairwise Welch's t-test. f. Representative bright field images of Numb-KD *vs.* Ctr-KD RT112 organoids treated as in ‘e’. Red arrowheads point to invasive protrusions. Bar, 50 µm. g. Transwell Matrigel invasion assay of Ctr-KD and Numb-KD RT112 cells treated for 24 h with VP (100 nM), C3 (3 µg/mL), Y-27632 (10 µM) or vehicle. Graph shows the average number of invading cells/field in each condition expressed as the mean ± SEM of 8 fields from two independent experiments. ****p<0.0001, ***, p<0.001; *, p<0.05; ns, not significant, relative to matching controls by FDR-adjusted pairwise Welch's t-test.
